# Using Major Depression Polygenic Risk Scores to Explore the Depressive Symptom Continuum

**DOI:** 10.1101/2020.02.25.962704

**Authors:** Bradley S Jermy, Saskia P Hagenaars, Kylie P Glanville, Jonathan RI Coleman, David M Howard, Gerome Breen, Evangelos Vassos, Cathryn M Lewis

## Abstract

**Background:** It is not clear whether major depression (MD) is a categorical disorder or if depressive symptoms exist on a continuum based on severity. Observational studies comparing sub-threshold and clinical depression suggest MD is continuous, but many do not explore the full continuum and are yet to consider genetics as a risk factor. This study sought to understand if polygenic risk for MD could provide insight into the continuous nature of MD.

**Methods:** Factor analysis on symptom-level data from the UK Biobank (N=148,957) was used to derive continuous depression phenotypes which were tested for association with polygenic risk scores for a categorical definition of MD (N=119,692).

**Results:** Confirmatory factor analysis showed a five-factor hierarchical model, incorporating 15 of the original 18 items, produced good fit to the observed covariance matrix (CFI = 0.992, TLI = 0.99, RMSEA = 0.038, SRMR = 0.031). MD PRS associated with each factor score (standardised ß range: 0.057 – 0.064) and the association remained when the sample was stratified into case- and control-only subsets. The case-only subset had an increased association compared to controls for all factors, shown via a significant interaction between lifetime MD diagnosis and MD PRS (p-value range: 2.28×10^−3^ - 4.56×10^−7^).

**Conclusions:** An association between MD PRS and a continuous phenotype of depressive symptoms in case- and control-only subsets provides support against a purely categorical phenotype; indicating further insights into MD can be obtained when this within-group variation is considered. The stronger association within cases suggests this variation may be of particular importance.

## Introduction

Major Depression (MD) is a common psychiatric disorder that affects more than 300 million people worldwide (World Health Organization, 2017). The Diagnostic and Statistical Manual of Mental Health Disorders-5 (DSM-5) (American Psychiatric Association, 2013) and the International Classification of Disease 11 (ICD-11) (World Health Organization, 2018) implicitly assume a categorical model for MD, whereby a collection of symptoms reflects a common dysfunction not present in healthy individuals (Helzer, Kraemer & Krueger, 2006; Kraemer, Noda & O’Hara, 2004). An alternative view is that MD is dimensional, existing along a continuum. Under this assumption, diagnostic boundaries using DSM-5 and ICD-11 criteria represent an arbitrary threshold along the distribution to partition ‘affected’ from ‘unaffected’ individuals (Vares, Salum, Spanemberg, Caldieraro & Fleck, 2015). If MD is dimensional, dichotomisation will reduce the power of a study to characterise the phenotype. Simulations showed that categorising a right skewed phenotype according to a clinical cut-off reduced the power to detect a genetic effect explaining 2% of the variance from 90.9% to 9.2% (van der Sluis, Posthuma, Nivard, Verhage & Dolan, 2013).

To compare dimensional and categorical models, studies have focused on clinical correlates between sub-threshold and clinical MD, arguing that a linear trend between the two groups supports a dimensional phenotype (reviewed in Solomon, Hagga & Arnow, 2001; Rodríguez, Nuevo, Chatterji & Ayuso-Mateos, 2012). These findings support a dimensional classification of MD, indicating an association between subthreshold MD and increased risk for disability, impairment, comorbidities and health care use (Hybels, Blazer & Pieper, 2001; Rucci et al., 2003; Cuijpers, de Graaf & van Dorsselaer, 2004) and a linear trend between MD severity and risk of a future episode (Kessler, Zhao, Blazer & Swartz, 1997; Kendler & Gardner, 1998).

There are three key limitations in the literature. First, by grouping individuals into sub-threshold and clinical MD, the full continuum of the phenotype is not explored. Second, grouping individuals in this way fails to account for symptomatic heterogeneity. According to the DSM-5, one of two ‘core’ symptoms - low mood or anhedonia – in combination with any other four symptoms meets the criteria for a MD diagnosis (American Psychiatric Association, 2013). This gives rise to a possible 227 symptom profiles, with evidence that many of these profiles present clinically (Olbert, Gala & Tupler, 2014; Zimmerman, Ellison, Young, Chelminski & Dalrymple, 2015). Finally, the symptoms assessed all follow DSM criteria for MD. Restricting the analysis in this way omits potentially important information, for example, it is known MD is highly comorbid with other disorders including generalised anxiety disorder (GAD) (Moffitt et al., 2007).

Previous studies have rarely compared dimensional and categorical phenotypes of genetic risk. A meta-analysis of twin studies involving 21,000 individuals estimated the heritability of MD to be 37% (95% CI: 31%–42%) (Sullivan, Neale & Kendler, 2000). A recent genome-wide association study (GWAS) using a broad definition of MD have identified 102 independent loci (Howard et al., 2019). These studies confirm a polygenic architecture for MD, where genetic predisposition comprises many common genetic variants of small effect that additively increase risk (Sullivan, Daly & O’Donovan, 2012). GWAS results can be used to determine a single measure of genetic liability from common variants to MD, the polygenic risk score (PRS), calculated by summing the number of risk alleles carried by an individual, weighted by their effect size (Wray et al., 2014). How PRS associates with various characterisations of MD can be used to inform our nosological understanding of the disorder.

This study aimed to assess evidence for a dimensional vs categorical basis of MD using symptom and genetic data. Exploratory and confirmatory factor analysis were used to construct multiple dimensional phenotypes for MD within the UK Biobank, a volunteer-based, national health resource. These phenotypes were tested for association with MD PRS - calculated according to a categorical definition. Individuals were stratified by self-reported MD status to explore how the association differed between MD cases and controls.

## Methods

### Participants

Participants were from the UK Biobank, a national health resource of 502,655 individuals aged between 37 and 73 at the time of recruitment (2006 - 2010) (Bycroft et al., 2018). In 2016, participants were offered an online Mental Health Questionnaire (MHQ) which 157,336 participants voluntarily completed (Davis et al., 2020).

### Item Selection for the Dimensional Phenotypes

Eighteen items were selected from the MHQ to construct the dimensional phenotypes for MD. This included all items from the Patient Health Questionnaire 9 (PHQ-9) - a measure of depressive symptoms over the last two weeks which correspond to the DSM criteria for MD (Kroenke, Spitzer & Williams, 2001), all items from the General Anxiety Disorder 7 (GAD-7) (Spitzer, Kroenke & Williams, 2006), relating to symptoms of anxiety over the last two weeks, and two items from the subjective well-being questionnaire, on general happiness and how meaningful they believe their life to be. For a detailed list of these items, see Table S1.

148,957 MHQ participants provided a response to all items above. This sample was randomly and equally split into ‘training’ (N=74,478) and ‘test’ (N=74,479) sub-samples. The training sample was used to develop a factor model with good fit, and the test sample was used to internally validate this model using exploratory and confirmatory factor analysis.

### Exploratory factor analysis

Polychoric correlations were computed for the 18 ordinal items (Carroll, 1961) and ordinal alpha (Gadermann, Guhn & Zumbo, 2012) Keiser Meyer-Olkin (KMO) (Kaiser, 1974) and Bartlett’s test of sphericity (Bartlett, 1950) were computed. Factor analysis was considered an acceptable method if ordinal alpha and KMO were greater than 0.80, and Bartlett’s test with p<0.05 in line with previous guidelines (Table S2) (Gadermann et al., 2012; Beavers et al., 2013).

Exploratory Factor Analysis (EFA) was performed using the weighted least squares method and factors were allowed to rotate. Parallel analysis (Horn, 1965) and Velicer’s Minimum Average Partial test (Velicer, 1976) were used to produce an upper and lower bound for the appropriate number of factors respectively. Based on these bounds, factor models were fit iteratively and compared using the criteria: Tucker Lewis Index (TLI) ≥ 0.95, Root Mean Square Error Approximation (RMSEA) ≤ 0.05 and a smaller Bayesian Information Criteria (BIC) relative to other models. The model with the best fit was retained for further testing.

Items were removed according to a series of post-hoc tests using Thurstone’s analytical method for simple structure criteria (Thurstone, 1947) to attain good model fit. Where multiple models demonstrated good fit, the model retaining the largest number of items was chosen. All analysis steps, for EFA were performed using the psych package in R 3.4.1 (Revelle, 2017).

### Confirmatory factor analysis

To internally validate the EFA-derived model, confirmatory factor analysis (CFA) was performed in the ‘test’ sub-sample using the lavaan package in R 3.4.1 (Rosseel, 2012). Factor loadings greater than 0.3 from the EFA model were used to specify the relationships between latent variables and items within the CFA model. As in EFA, factors were allowed to rotate. In addition to the model fit metrics used in EFA, the comparative fit index (CFI) and standardized root mean square residual (SRMR) were calculated. Models with CFI ≥ 0.95 and SRMR ≤ 0.05 were considered a good fit.

### Deriving the Phenotype - Factor Score Calculation

Following confirmation of the proposed model, the selected model was fit to the full sample using CFA, and factor scores were computed for each factor (Figure 1).

**Fig 1.**
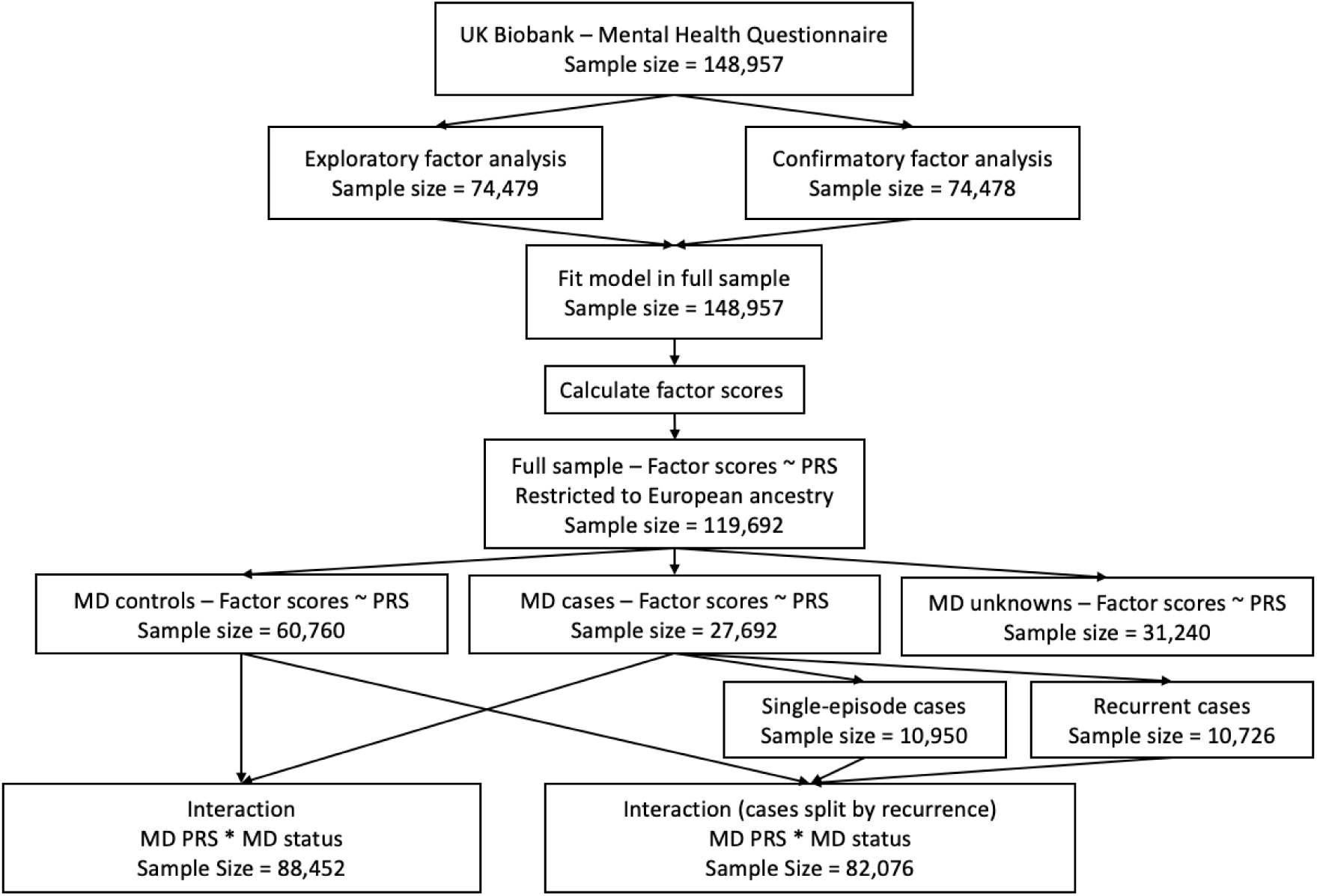
Flow chart displaying the methodology of the study along with the sample sizes for each section.

### Polygenic Risk Score Calculation

MD PRS were constructed using PRSice v2 (Euesden, Lewis & O’Reilly, 2014) in unrelated individuals of European ancestry (N = 119,692) using genotype data and quality control procedures previously described (Supplementary Methods) (Bycroft et al., 2018; Coleman et al., 2020). Summary statistics from Wray et al. (2018) with 23andMe and UK Biobank samples removed (N_cases_=45,591, N_controls_= 97,674) were used as the base dataset. To account for linkage disequilibrium, clumping was performed so that single nucleotide polymorphisms (SNPs) had an *r*^2^ < 0.1 and a 250kb window from other SNPs. MD PRS were then calculated across 11 p-value thresholds (p < 5 × 10^−8^, p < 1 × 10^−5^, p < 0.001, p < 0.01, p < 0.05, p < 0.1, p < 0.2, p < 0.3, p < 0.4, p < 0.5, p < 1). PRS for height were used as a negative control, and were calculated in the same way as MD PRS, using summary statistics from Wood et al. (2014) (N=253,228) as the base dataset.

### Association Testing

Factor scores and MD PRS were standardised, and linear regression was performed using the 119,692 MHQ participants. Each factor score was regressed on MD and height PRS with the first 6 genetic principal components, genotyping batch and assessment centre fitted as covariates.

### Stratification by Major Depression diagnosis

MD cases and controls were identified using the Composite International Diagnostic Interview - Short Form (CIDI-SF), a structured self-report questionnaire focusing on depressive symptoms during an individual’s worst episode of depression (Supplementary Methods) (Kessler, Andrews, Mroczek, Ustun & Wittchen, 1998). Individuals who did not complete the CIDI-SF, or who were removed due to the exclusion criteria, were assigned an ‘unknown’ status, resulting in 60,760 controls, 27,692 cases and 31,240 unknowns.

Participants were stratified into cases and controls and the same set of linear regressions repeated in each group. To test for group differences, cases and controls were combined (N=88,452) and interaction terms for MD PRS and case-control status as well as for MD PRS and all covariates were placed into the linear regression model. Cases were further stratified into single episode and recurrent cases with recurrence defined as having self-reported 2 or more lifetime depressive episodes and the interaction test was re-performed. In all interaction tests, controls were set as the reference for MD status.

As many of the tests are highly correlated, a method proposed by Nyholt (2004) was used to correct for multiple testing. The ‘poolR’ package (Cinar & Viechtbauer, 2016) in R 3.6.1 showed the effective number of independent tests for the factor scores and MD PRS was 14, giving a Bonferroni corrected α<0.0036 (0.05/14).

## Results

### Exploratory Factor Analysis

Initially, a six-factor model produced the best fit for the 18 items from the PHQ-9, GAD-7 and the two subjective well-being questions (Table S3). Model fit statistics did not surpass pre-defined thresholds (TLI = 0.973, RMSEA = 0.054), suggesting either poor discrimination or a lack of shared variance for particular items.

To improve model fit, items were removed according to a series of post-hoc tests based on the loadings from this six-factor model (Table S4) (Thurstone, 1947). The following criteria produced a model with a good fit to the data that retained the greatest number of items: remove items with no factor loadings above 0.4, remove items with loadings to multiple factors above 0.3, retain MD symptoms from the PHQ-9 that would otherwise qualify for removal except for an item for feelings of inadequacy (test 6; Table S4). These criteria retained 15 of the original 18 items and a five-factor model produced a model with good fit (TLI = 0.981, RMSEA = 0.048). Items relating to trouble relaxing, irritability and feelings of inadequacy were excluded.

### Confirmatory Factor Analysis

In contrast to EFA which allows all items to load on all factors, CFA specifies the relationship between items and factors directly and provides a more stringent test for the proposed model (Thompson, 2004). CFA within the test sub-sample (N = 74,479) confirmed a good fit to the observed covariance matrix (CFI = 0.995, TLI = 0.993, RMSEA = 0.033, SRMR = 0.025). As correlations between the five factors were moderate to high (r = 0.574 – 0.848), a second order latent variable was included in the model; the model fit remained within the pre-specified thresholds (CFI = 0.992, TLI = 0.99, RMSEA = 0.038, SRMR = 0.031). Fit statistics did not change when fitting the model into the full sample (N = 148,957) using CFA (Table S5).

In the final model, the five first-order factors reflected feelings of anxiety, psychomotor-cognitive impairment, neurovegetative states, mood and subjective well-being. The second-order factor, representing the correlation between these factors, was termed the ‘Internalising’ factor (Figure 2). Calculated as the average squared factor loading, the variance explained by each first order factor ranged from 49% - 70%. The internalising factor explained 72% of the variance for all first-order factors. The high degree of variance explained supports the use of these factors as phenotypes for dimensional MD.

**Fig 2.**
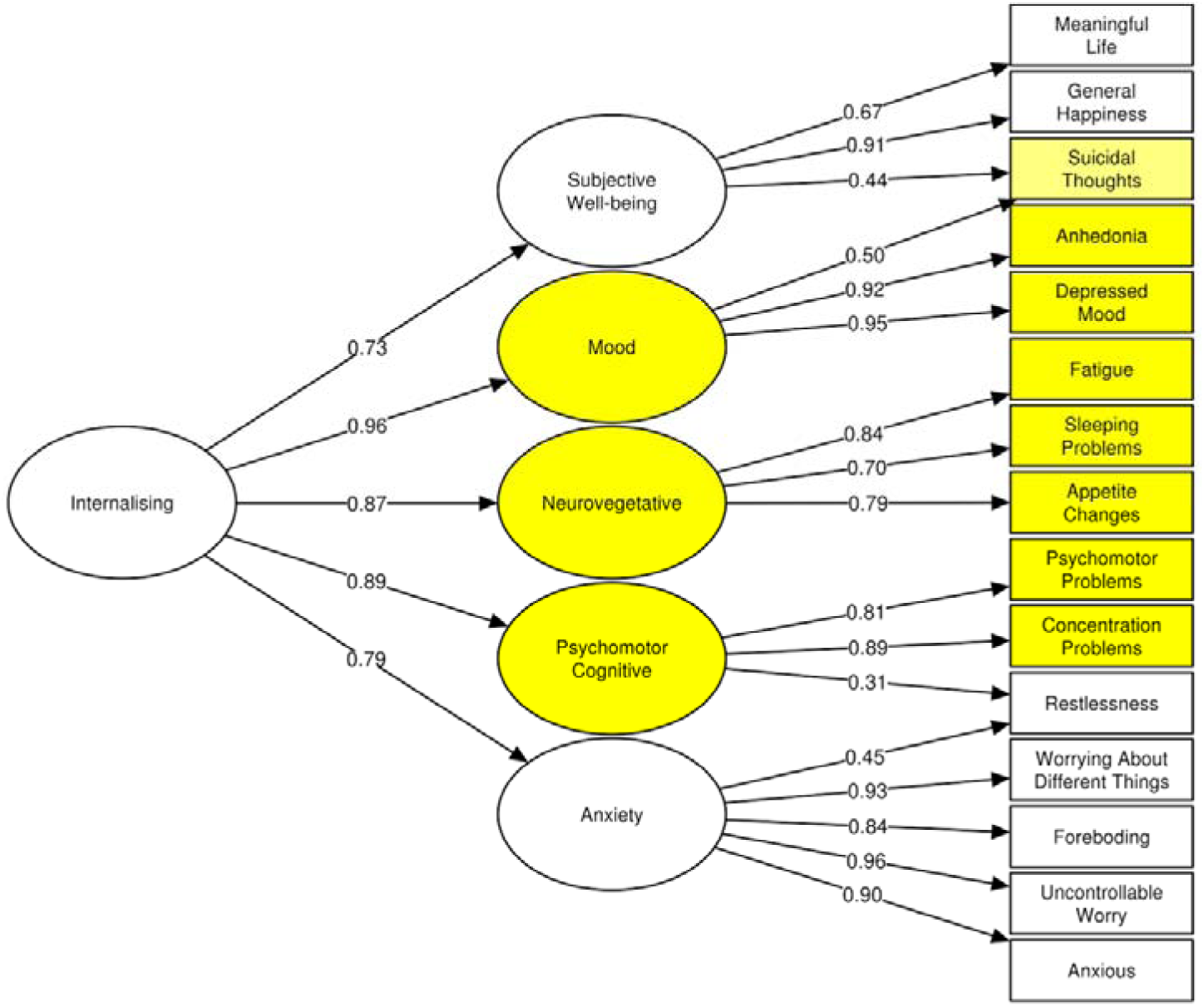
Factor model used to derive the dimensional phenotypes. As is customary in structural equation modelling graphs, circles are factors and squares are the self-reported symptoms. All symptoms shaded in yellow correspond to core MD symptoms with factors containing a majority of MD symptoms are also shaded. Arrows pointing from either one factor to a symptom or a factor to another factor represent the factor loadings.

### Major Depression Polygenic Risk Score Associates with an Individual’s Factor Score

MD PRS were associated with each of the factor scores (ß range: 0.056 – 0.064; p-value range: 2.57×10^−82^ - 1.89×10^−107^; MD PRS p-value threshold: p_T_ < 0.3) indicating their ability to differentiate between MD severity regardless of diagnostic status. While the effect sizes were similar across factors (Table S6), the two factors with the lowest association with MD PRS related to feelings of anxiety and subjective well-being (Figure 3). In contrast, height PRS was not associated with any factor score (p > 0.05) at any p-value threshold (Table S7).

**Fig 3.**
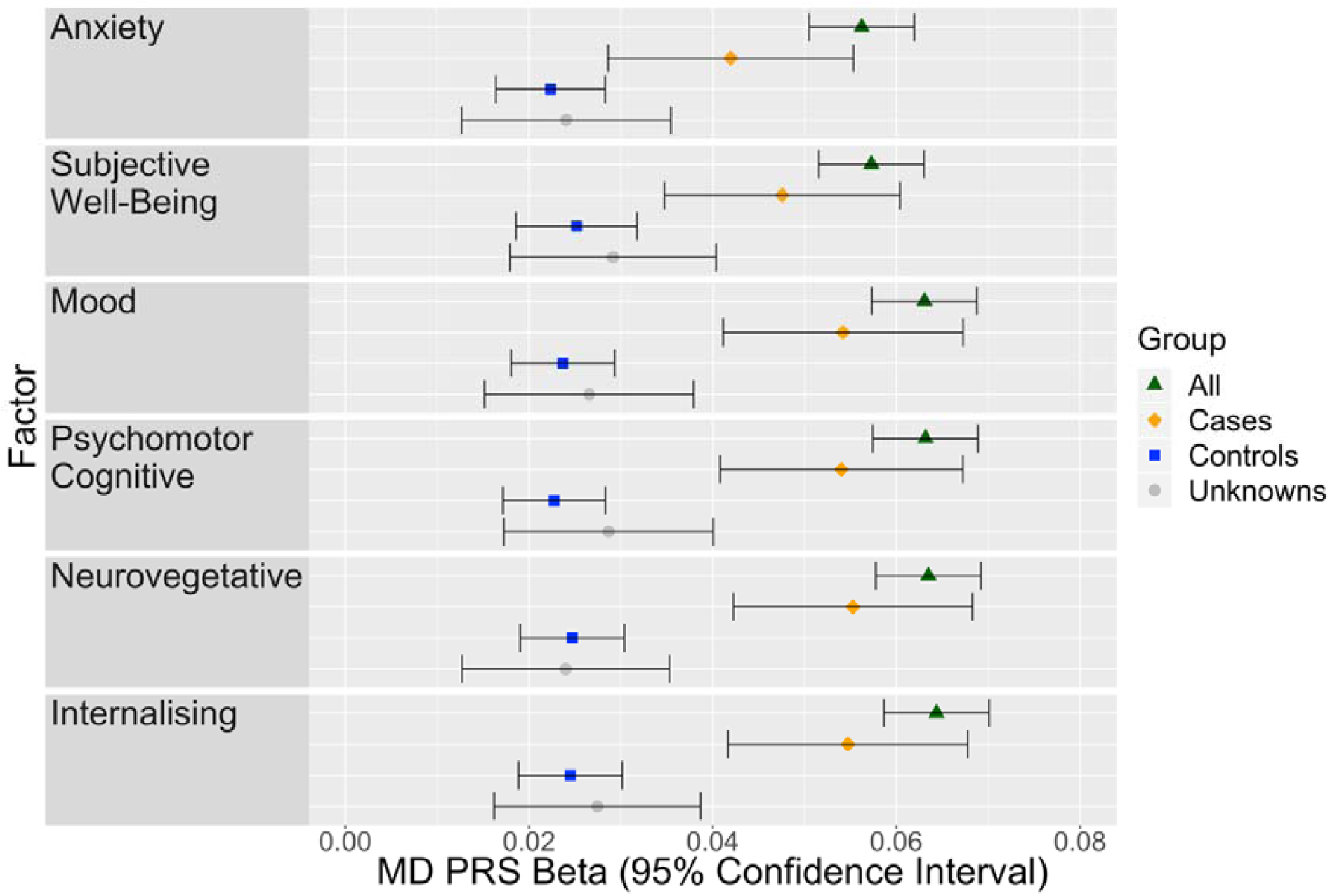
Association of MD PRS on each factor in the full sample and when stratified by MD Case/Control Status. The MD PRS used was calculated at the p-value threshold of p_T_ < 0.3. Both MD PRS and Factor scores were standardised to have a mean of 0 and variance of 1 using the full sample.

### Stratification by case/control status

MD PRS remained associated with factor scores following stratification into case- and control-only subgroups (controls: ß range: 0.022 – 0.025; p-value range: 1.89×10^−13^ - 1.43×10^−.17^; cases: ß range: 0.042 – 0.055; p-value range: 7.54×10^−10^ - 8.95×10^−17^; MD PRS p-value threshold: p_T_ < 0.3). Figure 3 shows the beta coefficients for MD PRS on each factor for each group (cases, controls and unknowns) (Tables S8-S10). The p-value thresholds of MD PRS with the largest effect size differed across factors between cases and controls (case threshold: p_T_ < 0.2; control threshold: p_T_ < 1) suggesting SNPs associated with MD contained greater signal for dimensions within cases whereas dimensions within controls required additional SNPs with weaker evidence for association in the base data.

MD PRS had an attenuated effect in controls relative to cases (Figure 3). To formally test for a differential genetic effect, factor scores were regressed on MD PRS and MD diagnostic status (case or control; N=88,452) with an interaction term. An interaction was detected for all factors (ß range: 0.020 – 0.031; p-value range: 2.28×10^−3^ - 4.56×10^−7^; MD PRS p-value threshold: p_T_ < 0.3; Table 1) indicating the genetic contribution is potentiated in cases relative to controls.

**Table 1:** Main and interaction effects of MD PRS and MD diagnostic status on an individual’s standardised factor score. The full sample has been subset to only include individuals with MD diagnostic status (*N*=27,692 cases; *N*=60,760 controls). Main effect models for MD PRS and MD diagnostic status include the relevant variable and covariates

| Factor | MD PRS (Main Effect) |  |  | MD diagnostic status (Main Effect) |  |  | MD PRS x MD diagnostic status |  |  |
| --- | --- | --- | --- | --- | --- | --- | --- | --- | --- |
| | Effect Size ($\beta$) | Std Error | P-value | Effect Size ($\beta$) | Std Error | P-value* | Effect Size ($\beta$) | Std Error | P-value |
| Anxiety | 0.061 | $3.28 \times 10^{-3}$ | $2.48 \times 10^{-77}$ | 0.891 | $6.34 \times 10^{-3}$ | $< 2 \times 10^{-16}$ | 0.020 | $6.43 \times 10^{-3}$ | $2.28 \times 10^{-3}$ |
| Psychomotor<br>Cognitive | 0.068 | $3.23 \times 10^{-3}$ | $4.58 \times 10^{-97}$ | 0.961 | $6.11 \times 10^{-3}$ | $< 2 \times 10^{-16}$ | 0.031 | $6.20 \times 10^{-3}$ | $4.56 \times 10^{-7}$ |
| Neurovegetative | 0.070 | $3.23 \times 10^{-3}$ | $3.13 \times 10^{-102}$ | 0.961 | $6.11 \times 10^{-3}$ | $< 2 \times 10^{-16}$ | 0.031 | $6.20 \times 10^{-3}$ | $8.32 \times 10^{-7}$ |
| Mood | 0.068 | $3.23 \times 10^{-3}$ | $8.07 \times 10^{-99}$ | 0.952 | $6.11 \times 10^{-3}$ | $< 2 \times 10^{-16}$ | 0.031 | $6.20 \times 10^{-3}$ | $8.48 \times 10^{-7}$ |
| Subjective Well-Being | 0.061 | $3.32 \times 10^{-3}$ | $1.13 \times 10^{-76}$ | 0.800 | $6.56 \times 10^{-3}$ | $< 2 \times 10^{-16}$ | 0.022 | $6.66 \times 10^{-3}$ | $7.66 \times 10^{-4}$ |
| Internalising | 0.070 | $3.24 \times 10^{-3}$ | $6.06 \times 10^{-102}$ | 0.973 | $6.12 \times 10^{-3}$ | $< 2 \times 10^{-16}$ | 0.030 | $6.21 \times 10^{-3}$ | $1.11 \times 10^{-6}$ |

\* P-value not possible to determine through `lm()` function in R as it is below the floor limit of the software. As such it is specified to simply be under a specific value given in the summary results from R ( $p < 2 \times 10^{-16}$ ).

Cases were stratified into single episode (N=10,590) and recurrent cases, reporting 2 or more lifetime depressive episodes (N=10,726). The interaction test was repeated with controls as the reference category. No interaction effect was detected between controls and single episode cases (p > 0.05) for any factors. A nominally significant interaction was found between controls and recurrent cases for all factors, except anxiety and subjective well-being. No results survived correction for multiple testing (Table 2).

**Table 2a:**
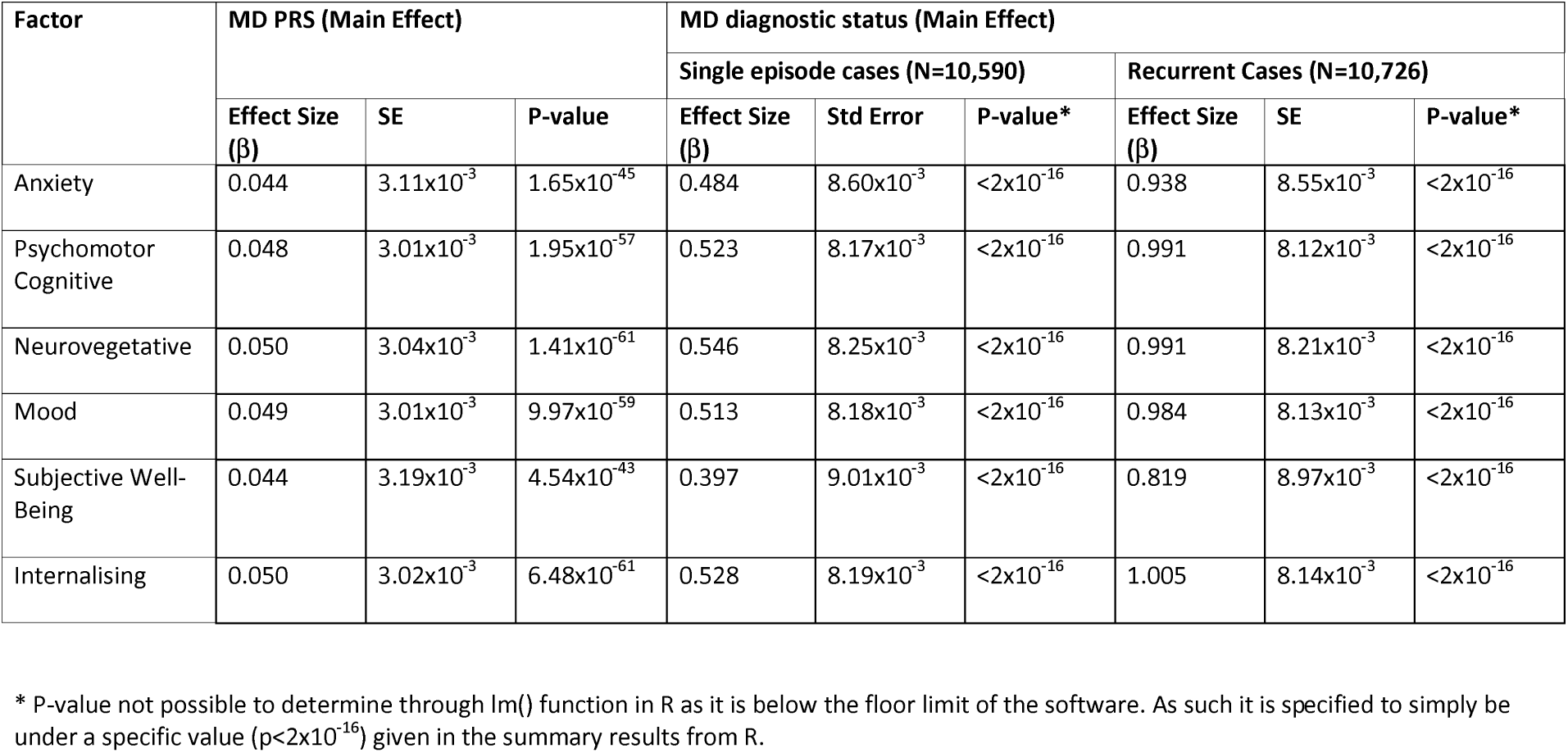
Main effects of MD PRS and MD diagnostic status split into single episode and recurrent cases. The full sample has been subset to only include individuals with MD diagnostic status who report the number of lifetime episodes of depression (*N*=10,590 single episode cases; *N*=10,726 recurrent cases; *N*=60,760 controls). Main effect models for MD PRS and MD diagnostic status include the relevant variable and covariates.

**Table 2b:** Interaction effects of MD PRS and MD diagnostic status split into single episode and recurrent cases. The full sample has been subset to only include individuals with MD diagnostic status who report the number of lifetime episodes of depression (*N*=10,590 single episode cases; *N*=10,726 recurrent cases; *N*=60,760 controls).

| Factor | MD PRS * MD Diagnostic status |  |  |  |  |  |
| --- | --- | --- | --- | --- | --- | --- |
| | Single episode cases ($N=10,590$) | | | Recurrent cases ($N=10,726$) | | |
| | Effect Size ($\beta$) | SE | P-value | Effect Size ($\beta$) | SE | P-value |
| Anxiety | -0.009 | $8.77 \times 10^{-3}$ | 0.288 | 0.008 | $8.74 \times 10^{-3}$ | 0.364 |
| Psychomotor<br>Cognitive | -0.005 | $8.33 \times 10^{-3}$ | 0.540 | 0.021 | $8.30 \times 10^{-3}$ | 0.013 |
| Neurovegetative | -0.008 | $8.41 \times 10^{-3}$ | 0.354 | 0.023 | $8.38 \times 10^{-3}$ | 0.006 |
| Mood | -0.006 | $8.34 \times 10^{-3}$ | 0.476 | 0.020 | $8.31 \times 10^{-3}$ | 0.016 |
| Subjective Well-Being | -0.008 | $9.20 \times 10^{-3}$ | 0.378 | 0.005 | $9.20 \times 10^{-3}$ | 0.572 |
| Internalising | -0.007 | $8.35 \times 10^{-3}$ | 0.420 | 0.019 | $8.32 \times 10^{-3}$ | 0.021 |

## Discussion

The aim of this study was to explore polygenic associations across the continuum of MD and test if this association differed between cases and controls. Using the UK Biobank, this study shows polygenic liability for a categorical MD phenotype (Wray et al., 2018) also associates with a dimensional model of depressive symptoms. Moreover, this finding holds in analyses stratified by case-control status. This suggests PRS contains information over and above the risk of becoming a case and may also be used to indicate severity across the continuum.

To account for symptom level heterogeneity within the dimensional phenotype, this study used factor analysis to derive a 5-factor hierarchical structure for MD. Previous studies investigating the latent structure of MD have produced multiple factor solutions. One and two factor models have been shown to produce the best fit for the PHQ-9, depending on the sample selected, i.e. case-only or population cohort (Elhai et al., 2012; Kocalevent, Hinz & Brähler, 2013). The 5-factor hierarchical model derived in this study has a high level of agreement for depressive symptoms with a model proposed by Kendler, Aggen & Neale (2013) in a population based sample of 7500 twins which showed three uncorrelated genetic factors best decompose the phenotypic variance in lifetime MD symptoms. Our model differed in two ways, firstly the symptom of feelings of worthlessness or excessive guilt was not included; and suicidal thoughts loaded on the mood factor, whereas it loaded on the psychomotor/cognitive factor in the model proposed by Kendler et al., (2013). Our study may therefore be considered a quasi-replication that used phenotypic rather than genetic covariance, in a substantially larger dataset (N=119,692), with current in contrast to lifetime symptoms, to provide support for the multidimensionality of MD.

Multidimensionality also indicates a deeper level of complexity in MD that is left unexplored when using either sum-scores of symptoms or a case-control design. A logical extension would be to use this multidimensionality to determine which genetic variants influence each factor, however, the high loadings of the 5 first order factors onto the second order ‘internalising’ factor suggests a significant proportion of this complexity is shared. As a result, it is likely that larger sample sizes will be required to identify genetic variants specific to a given factor.

The separation between symptoms of MD, anxiety and subjective well-being is noteworthy. The GAD-7 has previously been shown to possess a unidimensional factor structure (Löwe et al., 2008) distinct from symptoms of MD (Spitzer et al., 2006). Compared with the ‘core MD factors’, an interaction between controls and recurrent MD with MD PRS was not detected for the factors relating to anxiety and subjective well-being at the level of nominal significance. This may suggest that although these symptom dimensions contain a highly pleiotropic genetic component (Purves et al., 2020), a diagnosis of MD contains a degree of specificity which reflects the structure suggested by the DSM and ICD. However, as no factors survived correction for multiple testing for the recurrent case interaction, this conclusion warrants further investigation.

When stratified into cases and controls for MD, an attenuated association between MD PRS and the continuous phenotypes was evident for controls relative to cases. This is, perhaps, not surprising as controls were screened for the presence of any psychiatric disorders and high levels of current depressive symptoms (PHQ-9 sum-score < 14). As such, compared to cases, controls are expected to form a more homogenous group of healthy individuals, limiting the power for MD PRS to associate with the phenotype. Nevertheless, we still detected a significant association, suggesting that even in what would typically be considered a healthy group, MD PRS can still differentiate the subtle differences of the continuous phenotype.

In contrast, the effect size within cases was similar to that of the entire sample, indicating cases contain the majority of the signal for the dimensions. This finding has important implications under the assumption of a purely dimensional phenotype as it suggests ignoring the variation within cases, also ignores a substantial proportion of the association. The increased association within cases may be due to the increased variability within the group. Alternatively, questionnaires may be being interpreted differently between cases and controls for MD, perhaps due to a greater degree of familiarity with the questionnaires in cases. Familiarity may increase the validity of the responses as individuals are more ‘in tune’ with the symptoms, reducing measurement error and improving power for the study to detect an association. Importantly, the different associations between cases and controls does not provide evidence for a categorical phenotype for MD. It is possible to derive an interaction in this way by sampling from the tails of the distributions of a continuous variable.

Whilst the presence of an association within cases and controls for MD appears to contradict the purely categorical phenotype of MD, it does not exclude the possibility that MD is a categorical phenotype characterised by continuous variation within cases and controls. Two taxometric studies, designed specifically to detect the presence of such groups or ‘taxons’ provide support for this finding (Ruscio, Zimmerman, McGlinchey, Chelminski & Young, 2007; Ruscio, Brown & Ruscio, 2009). However, this field of research has consistently supported a dimensional classification, and the results appear to depend on the measurement instrument used, whether the symptoms were self-report or clinically ascertained and age of the sample (Hankin, Fraley, Lahey & Waldman, 2005; Liu, 2016).

## Limitations

This study has many strengths including large sample size from a volunteer-based, national health resource and its accounting for the heterogeneity inherent to MD, however, limitations remain. The items used to create the dimensional phenotypes were self-reported, increasing the risk of misclassification and sampling-bias. It has been shown that participants who responded to the MHQ have a higher level of education and fewer hospital diagnoses inclusive of mental disorders compared with other UK Biobank participants (Adams et al., 2019). This ‘healthier and wealthier’ bias may hamper our ability to appropriately represent cases at the most severe end of the spectrum. The same study showed that MHQ responders were more likely to have a family history of severe depression relative to non-responders (Adams et al., 2019). Similarly, the influence of personal interest in mental health could limit the generalizability of this sample to the general population.

This study assessed current depressive symptoms at a single time point, when the MHQ was completed. Studies investigating latent class trajectory of depressive symptoms have supported the dynamic nature of MD, with trajectories including persistently low, persistently high, increasing and decreasing symptoms (Kuchibhatla, Fillenbaum, Hybels & Blazer, 2012; Byers et al., 2012). Future studies should seek to confirm this model in longitudinal settings to test if it is robust to temporal invariance (Widamann, Ferrer & Conger, 2010). Furthermore, the MHQ was completed in one sitting, and the distinction between factors might reflect an artefact of time from the three questionnaires taken at different points in the MHQ (Davis et al., 2020).

## Conclusions

MD PRS supports a multi-dimensional model of MD, indicating that information is contained within cases and controls that would otherwise be omitted using a categorical phenotype. Much of this additional information is held within cases. Considering this in future study design may enhance the power to detect genetic associations, elevate our current aetiological understanding, and improve prediction through more accurate PRS.

## Acknowledgements

CML is funded by the Medical Research Council (N015746/1). SPH is funded by the Medical Research Council (MR/S0151132). DMH is supported by a Sir Henry Wellcome Postdoctoral Fellowship (Reference 213674/Z/18/Z) and a 2018 NARSAD Young Investigator Grant from the Brain & Behavior Research Foundation (Ref: 27404). This study represents independent research funded by the National Institute for Health Research (NIHR) Biomedical Research Centre at South London and Maudsley NHS Foundation Trust and King’s College London. The views expressed are those of the author(s) and not necessarily those of the NHS, the NIHR or the Department of Health and Social Care.

We thank participants and scientists involved in making the UK Biobank resource available (http://www.ukbiobank.ac.uk/). UK Biobank data used in this study were obtained under approved application 18177.

## References

Adams, M.J., Hill, W.D., Howard, D.M., Dashti, H.S., Davis, K.A.S., Campbell, A., …McIntosh, A.M. (2019). Factors associated with sharing e-mail information and mental health survey participation in large population cohorts. International journal of epidemiology. doi: 10.1093/ije/dyz134

American Psychiatric Association. (2013). Diagnostic and statistical manual of mental disorders (DSM-5^®^). American Psychiatric Publishing.

Bartlett, M.S. (1950). Tests of Significance in Factor Analysis. British Journal of statistical psychology, 3(2), 77–85. doi: 10.1111/j.2044-8317.1950.tb00285.x

Beavers, A.S., Lounsbury, J.W., Richards, J.K., Huck, S.W., Skolits, G.J., & Esquivel, S.L. (2013). Practical considerations for using exploratory factor analysis in educational research. Practical assessment, research & evaluation, 18(1), 6. Retrieved from: https://scholarworks.umass.edu/cgi/viewcontent.cgi?article=1303&context=pare

Bycroft, C., Freeman, C., Petkova, D., Band, G., Elliott, L.T., Sharp, K., … Marchini, J. (2018). The UK Biobank resource with deep phenotyping and genomic data. Nature, 562(7726), 203. doi: 10.1038/s41586-018-0579-z

Byers, A.L., Vittinghoff, E., Lui, L-Y., Hoang, T., Blazer, D.G., Covinsky, K.E.…, Yaffe, K. (2012). Twenty-Year Depressive Trajectories Among Older Women. Archives of general psychiatry, 69(10), 1073–9. doi: 10.1001/archgenpsychiatry.2012.43

Carroll, J.B. (1961). The nature of the data, or how to choose a correlation coefficient. Psychometrika, 26(4), 347–72. doi: 10.1007/BF02289768

Cinar, O., Viechtbauer, W. (2016). PoolR: Package for pooling the results from (dependent) tests. Retrieved from: https://rdrr.io/github/ozancinar/poolR/man/poolr-package.html

Coleman, J. R., Peyrot, W. J., Purves, K. L., Davis, K. A., Rayner, C., Choi, S. W., …Breen, G. (2020). Genome-wide gene-environment analyses of major depressive disorder and reported lifetime traumatic experiences in UK Biobank. Molecular psychiatry, 1–17. doi: 10.1038/s41380-019-0546-6

Cuijpers P, de Graaf R, & van Dorsselaer S. (2004). Minor depression: risk profiles, functional disability, health care use and risk of developing major depression. Journal of affective disorders, 79(1), 71–9. doi: 10.1016/S0165-0327(02)00348-8

Davis, K. A., Coleman, J. R., Adams, M., Allen, N., Breen, G., Cullen, B., …Hotopf, M. (2020). Mental health in UK Biobank–development, implementation and results from an online questionnaire completed by 157 366 participants: a reanalysis. BJPsych open, 6(2). doi: 10.1192/bjo.2019.100

Elhai, J.D., Contractor, A.A., Tamburrino, M., Fine TH, Prescott MR, Shirley E, …Calabrese, J.R. (2012). The factor structure of major depression symptoms: A test of four competing models using the Patient Health Questionnaire-9. Psychiatry research, 199(3), 169–73. doi: 10.1016/j.psychres.2012.05.018

Euesden, J., Lewis, C.M., & O’Reilly, P.F. (2014). PRSice: polygenic risk score software. Bioinformatics, 31(9), 1466–8. doi: 10.1093/bioinformatics/btu848

Gadermann, A. M., Guhn, M., Zumbo, & Bruno D. (2012). Estimating ordinal reliability for Likerttype and ordinal item response data: A conceptual, empirical, and practical guide. Practical assessment, research, and evaluation, 17(1), 3. Retrieved from: https://scholarworks.umass.edu/cgi/viewcontent.cgi?article=1247&context=pare

Hankin B.L., Fraley, R.C., Lahey, B.B., & Waldman, I.D. (2005). Is depression best viewed as a continuum or discrete category? A taxometric analysis of childhood and adolescent depression in a population-based sample. Journal of abnormal psychology, 114(1), 96–110 doi: 10.1037/0021-843X.114.1.96

Helzer, J.E., Kraemer, H.C., & Krueger, R.F. (2006). The feasibility and need for dimensional psychiatric diagnoses. Psychological medicine, 36(12), 1671–80. doi: 10.1017/S003329170600821X

Horn, J.L. (1965). A rationale and test for the number of factors in factor analysis. Psychometrika, 30(2), 179–85. doi: 10.1007/bf02289447

Howard, D.M., Adams, M.J., Clarke, T.K., Hafferty, J.D., Gibson, J., Shirali M, …McIntosh, A.M. (2019). Genome-wide meta-analysis of depression identifies 102 independent variants and highlights the importance of the prefrontal brain regions. Nature neuroscience, 22(3), 343. doi: 10.1038/s41593-018-0326-7

Hybels, C.F., Blazer, D.G., & Pieper, C.F. (2001). Toward a threshold for subthreshold depression: an analysis of correlates of depression by severity of symptoms using data from an elderly community sample. The Gerontologist, 41(3), 357–65. doi: 10.1093/geront/41.3.357

Kaiser, H.F. (1974). An index of factorial simplicity. Psychometrika, 39(1), 31–6. doi: 10.1007/BF02291575

Kendler, K.S., Aggen, S.H., & Neale, M.C. (2013). Evidence for multiple genetic factors underlying DSM-IV criteria for major depression. JAMA Psychiatry. 70(6), 599–607. doi: 10.1001/jamapsychiatry.2013.751.

Kendler, K.S., & Gardner, C.O. Jr. (1998). Boundaries of Major Depression: An Evaluation of DSM-IV Criteria. American journal of psychiatry, 155(2), 172–177. doi: 10.1176/ajp.155.2.172

Kessler, R.C., Andrews, G., Mroczek, D., Ustun, B., & Wittchen, H.U. (1998). The World Health Organization composite international diagnostic interview short-form (CIDI-SF). International journal of methods in psychiatric research, 7(4), 171–85. doi: 10.1002/mpr.47

Kessler, R.C., Zhao, S., Blazer, D.G., & Swartz, M. (1997). Prevalence, correlates, and course of minor depression and major depression in the National Comorbidity Survey. Journal of affective disorders, 45(1–2), 19–30. doi: 10.1016/s0165-0327(97)00056-6

Kocalevent, R-D., Hinz, A., & Brähler, E. (2013). Standardization of the depression screener Patient Health Questionnaire (PHQ-9) in the general population. General hospital psychiatry, 35(5), 551–5. doi: 10.1016/j.genhosppsych.2013.04.006

Kraemer, H.C., Noda, A., & O’Hara, R. (2004). Categorical versus dimensional approaches to diagnosis: methodological challenges. Journal of psychiatric research, 38(1), 17–25. doi: 10.1016/s0022-3956(03)00097-9

Kroenke, K., Spitzer, R.L., & Williams, J.B.W. (2001). The PHQ-9: validity of a brief depression severity measure. Journal of general internal medicine, 16(9), 606–13. doi: 10.1046/j.1525-1497.2001.016009606.x

Kuchibhatla, M.N., Fillenbaum, G.G., Hybels, C.F., & Blazer, D.G. (2012). Trajectory classes of depressive symptoms in a community sample of older adults. Acta psychiatrica scandinavica, 125(6), 492–501. doi: 10.1111/j.1600-0447.2011.01801.x

Liu, R.T. (2016). Taxometric evidence of a dimensional latent structure for depression in an epidemiological sample of children and adolescents. Psychological medicine, 46(6), 1265–75. doi: 10.1017/S0033291715002792

Löwe, B., Decker, O., Müller, S., Brähler, E., Schellberg, D., Herzog, W., & Herzberg, P.Y. (2008). Validation and standardization of the Generalized Anxiety Disorder Screener (GAD-7) in the general population. Medical care, 46(3), 266–74. doi: 10.1097/MLR.0b013e318160d093

Moffitt, T.E., Harrington, H., Caspi, A., Kim-Cohen, J., Goldberg, D., … Poulton, R. (2007). Depression and Generalized Anxiety Disorder: Cumulative and Sequential Comorbidity in a Birth Cohort Followed Prospectively to Age 32 Years. Archives of general psychiatry, 64(6), 651–60. doi: 10.1001/archpsyc.64.6.651

Nyholt, D.R. (2004). A simple correction for multiple testing for single-nucleotide polymorphisms in linkage disequilibrium with each other. The American journal of human genetics, 74(4), 765–9. doi: 10.1086/383251

Olbert, C.M., Gala, G.J., & Tupler, L.A. (2014). Quantifying heterogeneity attributable to polythetic diagnostic criteria: theoretical framework and empirical application. Journal of abnormal psychology, 123(2), 452–62. doi: 10.1037/a0036068

Purves, K. L., Coleman, J. R., Meier, S. M., Rayner, C., Davis, K. A., Cheesman, R., …Eley, T. C. (2019). A major role for common genetic variation in anxiety disorders. Molecular psychiatry, 1–12. doi: 10.1038/s41380-019-0559-1

Revelle, W. (2017). psych: Procedures for Psychological, Psychometric, and Personality Research. Retrieved from: https://CRAN.R-project.org/package=psych

Rodríguez, M.R., Nuevo, R., Chatterji, S., & Ayuso-Mateos, J.L. (2012). Definitions and factors associated with subthreshold depressive conditions: a systematic review. BMC Psychiatry, 12(1), 181. doi: 10.1186/1471-244X-12-181

Rosseel, Y. (2012). lavaan: An R Package for Structural Equation Modeling. Journal of statistical software, 48(1), 1–36. doi: 10.18637/jss.v048.i02

Rucci, P., Gherardi, S., Tansella, M., Piccinelli M, Berardi D, Bisoffi G, … Pini, S. (2003). Subthreshold psychiatric disorders in primary care: prevalence and associated characteristics. Journal of affective disorders, 76(1), 171–81. doi: 10.1016/s0165-0327(02)00087-3

Ruscio, J., Brown, T.A., & Ruscio, A.M. (2009). A Taxometric Investigation of DSM-IV Major Depression in a Large Outpatient Sample. Assessment, 16(2), 127–44. doi: 10.1177/1073191108330065

Ruscio, J., Zimmerman, M., McGlinchey, J.B., Chelminski, I., & Young, D. (2007). Diagnosing Major Depressive Disorder XI: A Taxometric Investigation of the Structure Underlying DSM-IV Symptoms. The Journal of nervous and mental disease, 195(1), 10–19. doi: 10.1097/01.nmd.0000252025.12014.c4

Solomon, A., Haaga, D.A., & Arnow, B.A. (2001). Is clinical depression distinct from subthreshold depressive symptoms? A review of the continuity issue in depression research. The Journal of nervous and mental disease, 189(8), 498–506. doi: 10.1097/00005053-200108000-00002

Spitzer, R.L., Kroenke, K., Williams, J.B.W., & Löwe, B. (2006). A brief measure for assessing generalized anxiety disorder: the GAD-7. Archives of internal medicine, 166(10), 1092–7. doi: 10.1001/archinte.166.10.1092

Sullivan, P.F., Daly, M.J., & O’Donovan, M. (2012). Genetic architectures of psychiatric disorders: the emerging picture and its implications. Nature reviews genetics, 13(8), 537–51. doi: 10.1038/nrg3240

Sullivan, P.F., Neale, M.C., & Kendler, K.S. (2000). Genetic Epidemiology of Major Depression: Review and Meta-Analysis. American journal of psychiatry, 157(10), 1552–62. doi: 10.1176/appi.ajp.157.10.1552

Thompson, B. (2004). Exploratory and confirmatory factor analysis: Understanding concepts and applications. Washington, DC, 10694.

Thurstone, L.L. (1947). Multiple factor analysis; a development and expansion of The Vectors of Mind. University of Chicago Press.

van der Sluis, S., Posthuma, D., Nivard, M.G., Verhage, M., & Dolan, C.V. (2013) Power in GWAS: lifting the curse of the clinical cut-off. Molecular psychiatry, 18(1), 2–3. doi: 10.1038/mp.2012.65

Vares, E.A., Salum, G.A., Spanemberg, L., Caldieraro, M.A., & Fleck, M.P. (2015). Depression Dimensions: Integrating Clinical Signs and Symptoms from the Perspectives of Clinicians and Patients. PLoS ONE, 10(8). doi: 10.1371/journal.pone.0136037

Velicer, W.F. (1976). Determining the number of components from the matrix of partial correlations. Psychometrika, 41(3), 321–7. doi: 10.1007/BF02293557

Widaman, K.F., Ferrer, E., & Conger, R.D. (2010). Factorial Invariance Within Longitudinal Structural Equation Models: Measuring the Same Construct Across Time. Child development perspectives, 4(1), 10–8. doi: 10.1111/j.1750-8606.2009.00110.x

Wood, A.R., Esko, T., Yang, J., Vedantam, S., Pers, T.H., Gustafsson S, … Frayling, T.M. (2014). Defining the role of common variation in the genomic and biological architecture of adult human height. Nature genetics, 46(11), 1173–86. doi: 10.1038/ng.3097

World Health Organization. (2017). Depression and other common mental disorders: global health estimates. Retrieved from: https://apps.who.int/iris/bitstream/handle/10665/254610/WHO-MSD-MER-2017.2-eng.pdf

World Health Organization. (2018). International classification of diseases for mortality and morbidity statistics (11^th^ Revision).

Wray, N.R., Lee, S.H., Mehta, D., Vinkhuyzen, A.A.E., Dudbridge, F., & Middeldorp, C.M. (2014). Research Review: Polygenic methods and their application to psychiatric traits. Journal of child psychology and psychiatry, 55(10), 1068–87. doi: 10.1111/jcpp.12295

Wray, N.R., Ripke, S., Mattheisen, M., Trzaskowski, M., Byrne, E.M., Abdellaoui, A., …the Major Depressive Disorder Working Group of the Psychiatric Genomics Consortium. (2018). Genome-wide association analyses identify 44 risk variants and refine the genetic architecture of major depression. Nature genetics, 50(5), 668. doi: 10.1038/s41588-018-0090-3

Zimmerman, M., Ellison, W., Young, D., Chelminski, I., & Dalrymple, K. (2015). How many different ways do patients meet the diagnostic criteria for major depressive disorder? Comprehensive psychiatry, 56, 29–34. doi: 10.1016/j.comppsych.2014.09.007

