## Supplementary Information for "Using Major Depression Polygenic Risk Scores to Explore the Depressive Symptom Continuum"

**Supplementary Methods** Genotyping quality control procedures and major depression case definition

**Table S1.** Items selected from UK Biobank for factor analysis model

**Table S2.** Data adequacy for factor analysis in sample used for exploratory factor analysis

**Table S2a.** Polychoric correlation matrix

**Table S2b.** Ordinal alpha, Keiser-Meyer Olkin and Bartletts Test results

**Table S3.** Exploratory factor analysis results across multiple factor solutions using all items

**Table S3a.** Exploratory factor analysis - model fit statistics

**Table S3b.** Exploratory factor analysis - factor loadings. Loadings > 0.3 highlighted in bold.

**Table S3c.** Exploratory factor analysis - factor correlations

**Table S4.** Exploratory factor analysis post-hoc tests

**Table S4a.** Exploratory factor analysis – post-hoc tests - model fit statistics

**Table S4b.** Test 6 - Exploratory factor analysis - factor loadings. Loadings > 0.3 highlighted in bold.

**Table S4c.** Test 6 - Exploratory factor analysis factor correlations

**Table S5.** Confirmatory factor analysis results – model fit statistics in full sample.

**Table S6.** Major depression polygenic risk score association test in full sample

**Table S7.** Height polygenic risk score association test in full sample

**Table S8.** Major depression polygenic risk score association test in case-only subsample

**Table S9.** Major depression polygenic risk score association test in control-only subsample

**Table S10.** Major depression polygenic risk score association test in unknown-only subsample

**Supplementary References**

**Supplementary Methods** Genotyping Quality Control Procedures and Major Depression Case Definition

**Genotyping Quality Control Procedures**

Polygenic Risk Score computation used the genotyped single nucleotide polymorphisms (SNPs). In addition to the centralised quality control procedures described in Bycroft et al., (2018), SNPs were restricted to minor allele frequency > 0.01 and Hardy-Weinberg equilibrium p > 10^-8^. Participants were removed if consent had been withdrawn, more than 2% of common variants were missing or if genetic sex did not equal self-reported sex (X-chromosome homozygosity (FX) < 0.9 for male self-report, FX > 0.5 for female self-report). European ancestry was taken as the largest cluster following 4-means clustering on the first two genetic principal components provided by the UK Biobank (Warren et al., 2017) and related participants with kinship coefficients r < 0.044 computed by KING (Manichaikul, Mychaleckyj, Rich, Daly, Sale & Chen, 2010) were removed. Removal of related participants used a ‘greedy’ algorithm to minimise reduction in sample size (i.e. retaining mother and father in the scenario the mother, father and child were all present). Flashpca2 was used to compute the genetic principal components in the participants of European ancestry for use as covariates in the linear regression portion of the analysis (Abraham, Qiu & Inouye, 2017).

**Major Depression Case Definition**

Cases were identified in line with the ‘Depression ever’ definition published by Davis et al., (2020). To qualify as a case, participants must have:

- Endorsed the two items corresponding to the cardinal symptoms of Major Depression (MD), depressed mood (field 20446) or anhedonia (field 20441).
- Endorsed 5 of a possible 8 symptoms corresponding to the MD DSM-5 criteria (fields: 20435, 20450, 20446, 20441, 20449, 20536, 20532, 20437). Psychomotor retardation/agitation is the DSM-5 symptom not included within the questionnaire.
- Responded ‘Most of the day’ or ‘All day long’ when asked how much of the day the symptoms presented during the episode (field 20436).
- Responded ‘Almost every day’ or ‘every day’ when asked how often they felt this way (field 20439).
- Responded ‘Somewhat’ or ‘A lot’ to a question relating to the level of impairment these symptoms induced (field 20440).

**Major Depression Control Definition**

Controls were defined in line with the definition published in Coleman et al., (2020). To qualify as a control, participants must not:

- Be identified as a case in the above definition.
- Self-report mental health problems diagnosed by a professional (field 20544).
- Self-report depression in a previous interview with a psychiatric nurse (field 20002).
- Meet criteria for depression or bipolar disorder (field 20126)
- Have a hospital inpatient ICD10 code (primary or secondary diagnosis) for mood disorder (F30 – F39) (fields 41202 – 41204).
- Report anti-depressant use at baseline (field 20003).
- Report current symptoms of depression according to the PHQ-9: less than 14 on summed response (where "not at all" = 1 and "nearly every day" = 4) to recent (fields: 20507, 20508, 20510, 20511, 20513, 20514, 20517, 20518, 20519)

**Exclusion Criteria**

To reduce misclassification in the definitions, cases and controls were removed if the following mental health problems were self-reported (field 20544 – Have you been diagnosed with one or more of the following mental health problems diagnosed by a professional?):

- Mania, hypomania, bipolar, manic-depression, schizophrenia or any other type of psychosis.

**Table S1*.*** Items selected from UK Biobank for factor analysis model

| **Field** | **Symptom Class** | **Symptom** | **Question** |
| --- | --- | --- | --- |
| 20510 | Depressive Symptoms | Depressed mood | Over the last 2 weeks, how often have you been bothered by any of the following problems? Feeling down, depressed, or hopeless |
| 20514 | Depressive Symptoms | Anhedonia | Over the last 2 weeks, how often have you been bothered by any of the following problems? Little interest or pleasure in doing things |
| 20511 | Depressive Symptoms | Appetite loss or gain | Over the last 2 weeks, how often have you been bothered by any of the following problems? Poor appetite or overeating |
| 20517 | Depressive Symptoms | Insomnia or hypersomnia | Over the last 2 weeks, how often have you been bothered by any of the following problems? Trouble falling or staying asleep, or sleeping too much |
| 20518 | Depressive Symptoms | Psychomotor agitation or retardation | Over the last 2 weeks, how often have you been bothered by any of the following problems? Moving or speaking so slowly that other people could have noticed? Or the opposite - being so fidgety or restless that you have been moving around a lot more than usual |
| 20519 | Depressive Symptoms | Fatigue or loss of energy | Over the last 2 weeks, how often have you been bothered by any of the following problems? Feeling tired or having little energy |
| 20507 | Depressive Symptoms | Feelings of worthlessness or excessive guilt | Over the last 2 weeks, how often have you been bothered by any of the following problems? Feeling bad about yourself or that you are a failure or have let yourself or your family down |
| 20508 | Depressive Symptoms | Impaired ability to think, concentrate | Over the last 2 weeks, how often have you been bothered by any of the following problems? Trouble concentrating on things, such as reading the newspaper or watching television |
| 20513 | Depressive Symptoms | Recurrent thoughts of death or suicide ideation, plan for committing suicide. | Over the last 2 weeks, how often have you been bothered by any of the following problems? Thoughts that you would be better off dead or of hurting yourself in some way |
| 20506 | Anxiety Symptoms | Nervous, anxious or on edge | Over the last 2 weeks, how often have you been bothered by any of the following problems? Feeling nervous, anxious or on edge |
| 20509 | Anxiety Symptoms | Uncontrollable worrying | Over the last 2 weeks, how often have you been bothered by any of the following problems? Not being able to stop or control worrying |
| 20515 | Anxiety Symptoms | Trouble relaxing | Over the last 2 weeks, how often have you been bothered by any of the following problems? Trouble relaxing |
| 20505 | Anxiety Symptoms | Irritable | Over the last 2 weeks, how often have you been bothered by any of the following problems? Becoming easily annoyed or irritable |
| 20520 | Anxiety Symptoms | Worrying about different things | Over the last 2 weeks, how often have you been bothered by any of the following problems? Worrying too much about different things |
| 20512 | Anxiety Symptoms | Foreboding | Over the last 2 weeks, how often have you been bothered by any of the following problems? Feeling afraid as if something awful might happen |
| 20516 | Anxiety Symptoms | Restlessness | Over the last 2 weeks, how often have you been bothered by any of the following problems? Being so restless that it is hard to sit still |
| 20458 | Happiness and subjective well-being | General Happiness | In general, how happy are you? |
| 20460 | Happiness and subjective well-being | Belief that own life is meaningful | To what extent do you feel your life to be meaningful? |

**Table S2.** Data adequacy for factor analysis

**Table S2a.** Polychoric correlation matrix

|  | **GH** | **ML** | **Irr** | **Anx** | **WG** | **Con** | **UW** | **DM** | **App** | **For** | **ST** | **Anh** | **TR** | **Res** | **Sle** | **Psy** | **Fat** | **WD** |
| --- | --- | --- | --- | --- | --- | --- | --- | --- | --- | --- | --- | --- | --- | --- | --- | --- | --- | --- |
| **General happiness (GH)** | 1 |  |  |  |  |  |  |  |  |  |  |  |  |  |  |  |  |  |
| **Belief that own life is meaningful (ML)** | 0.67 | 1 |  |  |  |  |  |  |  |  |  |  |  |  |  |  |  |  |
| **Irritable (Irr)** | 0.49 | 0.39 | 1 |  |  |  |  |  |  |  |  |  |  |  |  |  |  |  |
| **Anxious (Anx)** | 0.52 | 0.4 | 0.66 | 1 |  |  |  |  |  |  |  |  |  |  |  |  |  |  |
| **Worthlessness/**  **Excessive guilt (WG)** | 0.62 | 0.57 | 0.59 | 0.64 | 1 |  |  |  |  |  |  |  |  |  |  |  |  |  |
| **Impaired concentration (Con)** | 0.52 | 0.45 | 0.56 | 0.57 | 0.67 | 1 |  |  |  |  |  |  |  |  |  |  |  |  |
| **Uncontrollable worrying (UW)** | 0.55 | 0.42 | 0.67 | 0.86 | 0.67 | 0.59 | 1 |  |  |  |  |  |  |  |  |  |  |  |
| **Depressed Mood (DM)** | 0.68 | 0.54 | 0.62 | 0.68 | 0.78 | 0.68 | 0.7 | 1 |  |  |  |  |  |  |  |  |  |  |
| **Appetite loss/gain (App)** | 0.46 | 0.4 | 0.49 | 0.48 | 0.61 | 0.61 | 0.51 | 0.61 | 1 |  |  |  |  |  |  |  |  |  |
| **Foreboding (For)** | 0.5 | 0.4 | 0.62 | 0.76 | 0.63 | 0.55 | 0.79 | 0.63 | 0.48 | 1 |  |  |  |  |  |  |  |  |
| **Suicidal thoughts (ST)** | 0.68 | 0.64 | 0.55 | 0.58 | 0.76 | 0.64 | 0.62 | 0.77 | 0.55 | 0.6 | 1 |  |  |  |  |  |  |  |
| **Anhedonia (Anh)** | 0.66 | 0.54 | 0.59 | 0.61 | 0.71 | 0.7 | 0.63 | 0.87 | 0.64 | 0.57 | 0.71 | 1 |  |  |  |  |  |  |
| **Trouble relaxing (TR)** | 0.53 | 0.4 | 0.68 | 0.77 | 0.62 | 0.64 | 0.8 | 0.66 | 0.53 | 0.69 | 0.57 | 0.62 | 1 |  |  |  |  |  |
| **Restlessness (Res)** | 0.4 | 0.33 | 0.63 | 0.62 | 0.5 | 0.59 | 0.65 | 0.53 | 0.47 | 0.61 | 0.49 | 0.5 | 0.79 | 1 |  |  |  |  |
| **Insomnia/Hypersomnia (Sle)** | 0.38 | 0.31 | 0.43 | 0.44 | 0.47 | 0.52 | 0.49 | 0.53 | 0.52 | 0.42 | 0.46 | 0.52 | 0.53 | 0.44 | 1 |  |  |  |
| **Psychomotor retardation/agitation (Psy)** | 0.46 | 0.38 | 0.53 | 0.54 | 0.58 | 0.69 | 0.55 | 0.6 | 0.56 | 0.52 | 0.59 | 0.61 | 0.58 | 0.64 | 0.47 | 1 |  |  |
| **Fatigue/Loss of energy (Fat)** | 0.48 | 0.4 | 0.52 | 0.51 | 0.57 | 0.63 | 0.52 | 0.65 | 0.64 | 0.47 | 0.55 | 0.68 | 0.56 | 0.44 | 0.62 | 0.57 | 1 |  |
| **Worrying about different things (WD)** | 0.52 | 0.4 | 0.67 | 0.82 | 0.66 | 0.58 | 0.91 | 0.68 | 0.5 | 0.77 | 0.59 | 0.61 | 0.79 | 0.64 | 0.48 | 0.53 | 0.52 | 1 |

**Table S2b.** Ordinal alpha, Keiser-Meyer Olkin and Bartletts Test results in exploratory factor analysis sample.

| **Test** | **Statistic** |
| --- | --- |
| Ordinal alpha | 0.96 |
| Kaiser-Meyer Olkin test | 0.96 |
| Bartletts test | 1252520 (p < 0.05) |

**Table S3.** Exploratory factor analysis results across multiple factor solutions using all items

**Table S3a.** Exploratory factor analysis model fit statistics

|  | **Two factor model** | **Three factor model** | **Four factor model** | **Five factor model** | **Six factor**  **model** |
| --- | --- | --- | --- | --- | --- |
| **TLI** | 0.861 | 0.901 | 0.937 | 0.949 | 0.973 |
| **RMSEA** | 0.123 | 0.104 | 0.083 | 0.075 | 0.054 |
| **BIC** | 132879.9 | 81401.43 | 44034.88 | 29880.49 | 12667.12 |
| **Mean item complexity** | 1.1 | 1.3 | 1.4 | 1.5 | 1.6 |
| **Variance explained** | 0.64 | 0.68 | 0.71 | 0.73 | 0.75 |

**Table S3b.** Exploratory factor analysis factor loadings. Loadings > 0.3 highlighted in bold.

|  | **Factor 1** | **Factor 2** | **Factor 3** | **Factor 4** | **Factor 5** | **Factor 6** |
| --- | --- | --- | --- | --- | --- | --- |
| **General happiness** | 0.06 | 0.01 | **0.72** | 0.02 | 0.09 | -0.06 |
| **Belief that own life is meaningful** | -0.03 | 0 | **0.94** | 0 | -0.22 | 0.02 |
| **Irritable** | **0.35** | 0.07 | 0.02 | 0.28 | 0.13 | 0.04 |
| **Anxious/Nervous** | **0.83** | -0.03 | -0.01 | 0.04 | 0.07 | 0.02 |
| **Feelings of inadequacy** | **0.34** | 0.06 | 0.24 | -0.04 | 0.12 | **0.34** |
| **Impaired concentration** | 0.02 | **0.33** | 0.05 | 0.28 | 0.06 | **0.3** |
| **Uncontrollable worrying** | **0.96** | 0.02 | 0 | -0.01 | 0 | -0.01 |
| **Depressed Mood** | 0.06 | 0 | 0 | 0.01 | **0.94** | 0.01 |
| **Appetite loss/gain** | 0.03 | **0.55** | 0.02 | 0.06 | 0.04 | 0.19 |
| **Foreboding** | **0.78** | -0.06 | 0.02 | 0.06 | -0.05 | 0.18 |
| **Suicidal thoughts** | 0.19 | 0 | **0.5** | 0.02 | 0.04 | **0.32** |
| **Anhedonia** | -0.02 | 0.26 | 0.13 | 0.04 | **0.55** | 0.05 |
| **Trouble relaxing** | **0.42** | 0.13 | 0.04 | **0.47** | 0.04 | -0.12 |
| **Restlessness** | 0.04 | -0.05 | -0.02 | **0.95** | -0.01 | 0.03 |
| **Insomnia/Hypersomnia** | 0.08 | **0.65** | 0 | 0.07 | -0.05 | -0.02 |
| **Psychomotor retardation/agitation** | -0.02 | 0.23 | -0.01 | **0.42** | -0.01 | **0.38** |
| **Fatigue/Loss of energy** | 0.01 | **0.87** | -0.01 | -0.05 | 0.04 | 0.01 |
| **Worrying about different things** | **0.93** | 0.04 | -0.01 | 0 | -0.01 | -0.01 |

**Table S3c.** Exploratory factor analysis factor correlations

|  | **Factor 1** | **Factor 2** | **Factor 3** | **Factor 4** | **Factor 5** | **Factor 6** |
| --- | --- | --- | --- | --- | --- | --- |
| **Factor 1** | 1 |  |  |  |  |  |
| **Factor 2** | 0.63 | 1 |  |  |  |  |
| **Factor 3** | 0.57 | 0.57 | 1 |  |  |  |
| **Factor 4** | 0.71 | 0.59 | 0.44 | 1 |  |  |
| **Factor 5** | 0.69 | 0.71 | 0.74 | 0.53 | 1 |  |
| **Factor 6** | 0.37 | 0.5 | 0.51 | 0.35 | 0.59 | 1 |

**Table S4.** Exploratory factor analysis post-hoc tests

**Table S4a.** Exploratory factor analysis – post-hoc tests – model fit statistics

| **Test 1:** Any items from the EFA reference model with cross-loadings above 0.3 in two instances, or items with no single loading above 0.3 to be removed | | | | | | | | | |
| --- | --- | --- | --- | --- | --- | --- | --- | --- | --- |
| **EFA fit statistics** | **Two factors** | **Three factors** | | **Four factors** | | **Five factors** | | **Six factors** | |
| TLI | 0.909 | 0.937 | | 0.964 | | 0.836 | | Did not test | |
| RMSEA | 0.112 | 0.093 | | 0.07 | | 0.151 | | Did not test | |
| BIC | 49390.09 | 26907.34 | | 11543.65 | | 38938.41 | | Did not test | |
| Mean item complexity | 1.1 | 1.2 | | 1.1 | | 1.2 | | Did not test | |
| Variance explained | 0.63 | 0.68 | | 0.71 | | 0.72 | | Did not test | |
| **Test 2:** Any items from the EFA reference model with cross-loadings above 0.4 in two instances, or items with no single loading above 0.3 to be removed | | | | | | | | | |
| **EFA fit statistics** | **Two factors** | **Three factors** | | **Four factors** | | **Five factors** | | **Six factors** | |
| TLI | 0.878 | 0.918 | | 0.944 | | Did not test | | Did not test | |
| RMSEA | 0.117 | 0.096 | | 0.079 | | Did not test | | Did not test | |
| BIC | 103916.4 | 59332.32 | | 34067.37 | | Did not test | | Did not test | |
| Mean item complexity | 1.1 | 1.3 | | 1.4 | | Did not test | | Did not test | |
| Variance explained | 0.63 | 0.68 | | 0.7 | | Did not test | | Did not test | |
| **Test 3:** Any items from the EFA reference model with cross-loadings above 0.3 in three instances, or items with no single loading above 0.4 to be removed | | | | | | | | | |
| **EFA fit statistics** | **Two factors** | **Three factors** | | **Four factors** | | **Five factors** | | **Six factors** | |
| TLI | 0.856 | 0.887 | | 0.933 | | 0.966 | | Did not test | |
| RMSEA | 0.137 | 0.121 | | 0.093 | | 0.067 | | Did not test | |
| BIC | 105455.6 | 68439.97 | | 32557.67 | | 12826.94 | | Did not test | |
| Mean item complexity | 1.1 | 1.3 | | 1.3 | | 1.3 | | Did not test | |
| Variance explained | 0.64 | 0.69 | | 0.73 | | 0.75 | | Did not test | |
| **Test 4:** Any items from the EFA reference model with cross-loadings above 0.3 in two instances, or items with no single loading above 0.4 to be removed | | | | | | | | | |
| **EFA fit statistics** | **Two factors** | | **Three factors** | | **Four factors** | | **Five factors** | | **Six factors** |
| TLI | 0.912 | | 0.944 | | 0.981 | | Did not test | | Did not test |
| RMSEA | 0.115 | | 0.092 | | 0.054 | | Did not test | | Did not test |
| BIC | 42381.88 | | 20501.28 | | 5033.52 | | Did not test | | Did not test |
| Mean item complexity | 1 | | 1.2 | | 1.1 | | Did not test | | Did not test |
| Variance explained | 0.64 | | 0.69 | | 0.72 | | Did not test | | Did not test |
| **Test 5:** Using test 4 as a reference but retaining all items classified as depressive symptoms | | | | | | | | | |
| **EFA fit statistics** | **Two factors** | | **Three factors** | | **Four factors** | | **Five factors** | | **Six factors** |
| TLI | 0.876 | | 0.917 | | 0.944 | | N/A^a^ | | 0.905 |
| RMSEA | 0.121 | | 0.099 | | 0.081 | | N/A^a^ | | 0.106 |
| BIC | 97161.72 | | 54359.92 | | 30133.67 | | N/A^a^ | | 32602.4 |
| Mean item complexity | 1.1 | | 1.3 | | 1.4 | | N/A^a^ | | 1.4 |
| Variance explained | 0.64 | | 0.68 | | 0.71 | | N/A^a^ | | 0.75 |

a – Heywood case present in test

| **Test 6:** Using test 5 as a reference but removing the depressive symptom relating to feelings of worthlessness/excessive guilt | | | | | |
| --- | --- | --- | --- | --- | --- |
| **Model fit statistics** | **Two factors** | **Three factors** | **Four factors** | **Five factors** | **Six factors** |
| TLI | 0.875 | 0.918 | 0.949 | 0.981 | Did not test |
| RMSEA | 0.124 | 0.101 | 0.08 | 0.048 | Did not test |
| BIC | 87100.55 | 47181.25 | 23805.62 | 6562.9 | Did not test |
| Mean item complexity | 1.1 | 1.2 | 1.3 | 1.3 | Did not test |
| Variance explained | 0.63 | 0.68 | 0.71 | 0.74 | Did not test |

**Table S4b.** Test 6 - Exploratory factor analysis - factor loadings. Loadings > 0.3 highlighted in bold.

|  | **Factor 1** | **Factor 2** | **Factor 3** | **Factor 4** | **Factor 5** |
| --- | --- | --- | --- | --- | --- |
| **General happiness** | 0.06 | 0.01 | 0.25 | **0.58** | -0.02 |
| **Belief that own life is meaningful** | -0.06 | 0.01 | -0.03 | **0.86** | -0.01 |
| **Anxious/Nervous** | **0.8** | -0.01 | 0.09 | 0.01 | 0.04 |
| **Impaired concentration** | 0.03 | 0.25 | 0.17 | 0.06 | **0.44** |
| **Uncontrollable worrying** | **0.93** | 0.06 | 0.02 | 0.01 | -0.02 |
| **Depressed Mood** | 0.05 | 0.01 | **0.98** | -0.03 | -0.01 |
| **Appetite loss/gain** | 0 | **0.51** | 0.12 | 0.03 | 0.17 |
| **Foreboding** | **0.71** | -0.04 | 0.05 | 0.05 | 0.15 |
| **Suicidal thoughts** | 0.07 | 0.01 | **0.33** | **0.44** | 0.18 |
| **Anhedonia** | -0.03 | 0.25 | **0.62** | 0.09 | 0.06 |
| **Restlessness** | **0.43** | -0.01 | -0.09 | -0.04 | **0.55** |
| **Insomnia/Hypersomnia** | 0.13 | **0.62** | -0.06 | -0.01 | 0.06 |
| **Psychomotor retardation/agitation** | -0.01 | 0.08 | 0.11 | 0.02 | **0.74** |
| **Fatigue/Loss of energy** | 0 | **0.89** | 0.03 | -0.01 | -0.04 |
| **Worrying about different things** | **0.89** | 0.09 | 0 | 0.01 | -0.02 |

**Table S4c.** Test 6 - Exploratory factor analysis - factor correlations

|  | **Factor 1** | **Factor 2** | **Factor 3** | **Factor 4** | **Factor 5** |
| --- | --- | --- | --- | --- | --- |
| **Factor 1** | 1 |  |  |  |  |
| **Factor 2** | 0.68 | 1 |  |  |  |
| **Factor 3** | 0.65 | 0.49 | 1 |  |  |
| **Factor 4** | 0.6 | 0.59 | 0.44 | 1 |  |
| **Factor 5** | 0.71 | 0.58 | 0.5 | 0.66 | 1 |

**Table S5.** Confirmatory factor analysis - model fit statistics in full sample

| **Model Fit Test** | **Model Fit Statistic** |
| --- | --- |
| CFI | 0.992 |
| TLI | 0.99 |
| RMSEA | 0.038 |
| SRMR | 0.031 |

**Table S6.** Major depression polygenic risk score association test in full sample

| **Outcome: Anxiety Factor Score** | | | |
| --- | --- | --- | --- |
| **PRS Threshold** | **Beta** | **SE** | **P-val** |
| P_T_ < 5e.8 | 0.01172 | 0.00289 | 4.97E-05 |
| P_T_ < 1e.5 | 0.01573 | 0.00289 | 5.49E-08 |
| P_T_ < 0.001 | 0.02782 | 0.00289 | 6.88E-22 |
| P_T_ < 0.01 | 0.04307 | 0.00290 | 6.23E-50 |
| P_T_ < 0.05 | 0.04971 | 0.00290 | 1.10E-65 |
| P_T_ < 0.1 | 0.05426 | 0.00291 | 2.43E-77 |
| P_T_ < 0.2 | 0.05615 | 0.00292 | 3.44E-82 |
| P_T_ < 0.3 | 0.05622 | 0.00292 | 2.57E-82 |
| P_T_ < 0.4 | 0.05513 | 0.00292 | 3.37E-79 |
| P_T_ < 0.5 | 0.05504 | 0.00292 | 4.87E-79 |
| P_T_ < 1 | 0.05489 | 0.00292 | 1.55E-78 |
| **Outcome: Mood Factor Score** | | | |
| **PRS Threshold** | **Beta** | **SE** | **P-val** |
| P_T_ < 5e.8 | 0.01152 | 0.00289 | 6.65E-05 |
| P_T_ < 1e.5 | 0.01599 | 0.00289 | 3.23E-08 |
| P_T_ < 0.001 | 0.03039 | 0.00289 | 8.13E-26 |
| P_T_ < 0.01 | 0.04579 | 0.00290 | 3.13E-56 |
| P_T_ < 0.05 | 0.05547 | 0.00290 | 2.25E-81 |
| P_T_ < 0.1 | 0.06009 | 0.00291 | 1.74E-94 |
| P_T_ < 0.2 | 0.06285 | 0.00292 | 1.42E-102 |
| P_T_ < 0.3 | 0.06306 | 0.00292 | 3.86E-103 |
| P_T_ < 0.4 | 0.06186 | 0.00292 | 2.91E-99 |
| P_T_ < 0.5 | 0.06209 | 0.00292 | 4.06E-100 |
| P_T_ < 1 | 0.06227 | 0.00292 | 1.31E-100 |
| **Outcome: Neurovegetative Factor Score** | | | |
| **PRS Threshold** | **Beta** | **SE** | **P-val** |
| P_T_ < 5e.8 | 0.01168 | 0.00289 | 5.32E-05 |
| P_T_ < 1e.5 | 0.01305 | 0.00289 | 6.43E-06 |
| P_T_ < 0.001 | 0.02885 | 0.00289 | 1.98E-23 |
| P_T_ < 0.01 | 0.04565 | 0.00290 | 6.73E-56 |
| P_T_ < 0.05 | 0.05623 | 0.00290 | 1.43E-83 |
| P_T_ < 0.1 | 0.06128 | 0.00291 | 3.40E-98 |
| P_T_ < 0.2 | 0.06344 | 0.00292 | 1.73E-104 |
| P_T_ < 0.3 | 0.06349 | 0.00292 | 1.59E-104 |
| P_T_ < 0.4 | 0.06246 | 0.00292 | 3.62E-101 |
| P_T_ < 0.5 | 0.06274 | 0.00292 | 3.31E-102 |
| P_T_ < 1 | 0.06286 | 0.00292 | 1.69E-102 |

| **Outcome: Psychomotor Cognitive Factor Score** | | | |
| --- | --- | --- | --- |
| **PRS Threshold** | **Beta** | **SE** | **P-val** |
| P_T_ < 5e.8 | 0.01230 | 0.00289 | 2.07E-05 |
| P_T_ < 1e.5 | 0.01597 | 0.00289 | 3.41E-08 |
| P_T_ < 0.001 | 0.03135 | 0.00289 | 2.27E-27 |
| P_T_ < 0.01 | 0.04670 | 0.00290 | 2.04E-58 |
| P_T_ < 0.05 | 0.05628 | 0.00290 | 1.04E-83 |
| P_T_ < 0.1 | 0.06084 | 0.00291 | 8.66E-97 |
| P_T_ < 0.2 | 0.06289 | 0.00292 | 1.04E-102 |
| P_T_ < 0.3 | 0.06317 | 0.00292 | 1.70E-103 |
| P_T_ < 0.4 | 0.06215 | 0.00292 | 3.37E-100 |
| P_T_ < 0.5 | 0.06230 | 0.00292 | 8.89E-101 |
| P_T_ < 1 | 0.06239 | 0.00292 | 5.65E-101 |
| **Outcome: Subjective Well-Being Factor Score** | | | |
| **PRS Threshold** | **Beta** | **SE** | **P-val** |
| P_T_ < 5e.8 | 0.00684 | 0.00289 | 0.01777 |
| P_T_ < 1e.5 | 0.01563 | 0.00289 | 6.38E-08 |
| P_T_ < 0.001 | 0.02757 | 0.00289 | 1.46E-21 |
| P_T_ < 0.01 | 0.04102 | 0.00290 | 1.60E-45 |
| P_T_ < 0.05 | 0.04953 | 0.00290 | 2.56E-65 |
| P_T_ < 0.1 | 0.05339 | 0.00291 | 4.84E-75 |
| P_T_ < 0.2 | 0.05694 | 0.00292 | 1.30E-84 |
| P_T_ < 0.3 | 0.05728 | 0.00292 | 1.61E-85 |
| P_T_ < 0.4 | 0.05636 | 0.00292 | 8.13E-83 |
| P_T_ < 0.5 | 0.05663 | 0.00292 | 1.11E-83 |
| P_T_ < 1 | 0.05689 | 0.00292 | 2.26E-84 |
| **Outcome: Internalising Factor Score** | | | |
| **PRS Threshold** | **Beta** | **SE** | **P-val** |
| P_T_ < 5e.8 | 0.01178 | 0.00289 | 4.57E-05 |
| P_T_ < 1e.5 | 0.01605 | 0.00289 | 2.89E-08 |
| P_T_ < 0.001 | 0.03098 | 0.00289 | 9.22E-27 |
| P_T_ < 0.01 | 0.04698 | 0.00290 | 4.18E-59 |
| P_T_ < 0.05 | 0.05678 | 0.00290 | 3.45E-85 |
| P_T_ < 0.1 | 0.06156 | 0.00291 | 4.39E-99 |
| P_T_ < 0.2 | 0.06418 | 0.00292 | 6.70E-107 |
| P_T_ < 0.3 | 0.06438 | 0.00292 | 1.89E-107 |
| P_T_ < 0.4 | 0.06323 | 0.00292 | 1.22E-103 |
| P_T_ < 0.5 | 0.06343 | 0.00292 | 1.94E-104 |
| P_T_ < 1 | 0.06357 | 0.00292 | 8.47E-105 |

**Table S7.** Height polygenic risk score association test in full sample

| **Outcome: Anxiety Factor Score** | | | |
| --- | --- | --- | --- |
| **PRS Threshold** | **Beta** | **SE** | **P-val** |
| P_T_ < 5e.8 | -0.00145 | 0.00292 | 0.61938 |
| P_T_ < 1e.5 | 0.00053 | 0.00297 | 0.85803 |
| P_T_ < 0.001 | -0.00005 | 0.00311 | 0.98723 |
| P_T_ < 0.01 | -0.00042 | 0.00338 | 0.90169 |
| P_T_ < 0.05 | 0.00146 | 0.00375 | 0.69834 |
| P_T_ < 0.1 | 0.00131 | 0.00394 | 0.73887 |
| P_T_ < 0.2 | 0.00181 | 0.00410 | 0.65927 |
| P_T_ < 0.3 | 0.00371 | 0.00419 | 0.37595 |
| P_T_ < 0.4 | 0.00393 | 0.00425 | 0.35561 |
| P_T_ < 0.5 | 0.00373 | 0.00428 | 0.38339 |
| P_T_ < 1 | 0.00343 | 0.00431 | 0.42585 |
| **Outcome: Mood Factor Score** | | | |
| **PRS Threshold** | **Beta** | **SE** | **P-val** |
| P_T_ < 5e.8 | 0.00050 | 0.00292 | 0.86371 |
| P_T_ < 1e.5 | 0.00056 | 0.00297 | 0.85148 |
| P_T_ < 0.001 | 0.00097 | 0.00311 | 0.75556 |
| P_T_ < 0.01 | -0.00147 | 0.00338 | 0.66320 |
| P_T_ < 0.05 | -0.00033 | 0.00375 | 0.93014 |
| P_T_ < 0.1 | -0.00077 | 0.00394 | 0.84539 |
| P_T_ < 0.2 | -0.00041 | 0.00410 | 0.92015 |
| P_T_ < 0.3 | 0.00139 | 0.00419 | 0.74095 |
| P_T_ < 0.4 | 0.00146 | 0.00425 | 0.73130 |
| P_T_ < 0.5 | 0.00118 | 0.00428 | 0.78313 |
| P_T_ < 1 | 0.00090 | 0.00430 | 0.83480 |
| **Outcome: Neurovegetative Factor Score** | | | |
| **PRS Threshold** | **Beta** | **SE** | **P-val** |
| P_T_ < 5e.8 | 0.00159 | 0.00292 | 0.58629 |
| P_T_ < 1e.5 | 0.00094 | 0.00297 | 0.75058 |
| P_T_ < 0.001 | 0.00075 | 0.00311 | 0.80847 |
| P_T_ < 0.01 | -0.00197 | 0.00338 | 0.56086 |
| P_T_ < 0.05 | -0.00133 | 0.00375 | 0.72328 |
| P_T_ < 0.1 | -0.00235 | 0.00394 | 0.55187 |
| P_T_ < 0.2 | -0.00194 | 0.00410 | 0.63666 |
| P_T_ < 0.3 | -0.00015 | 0.00419 | 0.97163 |
| P_T_ < 0.4 | -0.00025 | 0.00425 | 0.95323 |
| P_T_ < 0.5 | -0.00046 | 0.00428 | 0.91457 |
| P_T_ < 1 | -0.00062 | 0.00430 | 0.88620 |

| **Outcome: Psychomotor Cognitive Factor Score** | | | |
| --- | --- | --- | --- |
| **PRS Threshold** | **Beta** | **SE** | **P-val** |
| P_T_ < 5e.8 | 0.00097 | 0.00292 | 0.73986 |
| P_T_ < 1e.5 | 0.00123 | 0.00297 | 0.67899 |
| P_T_ < 0.001 | 0.00115 | 0.00311 | 0.71211 |
| P_T_ < 0.01 | -0.00122 | 0.00338 | 0.71812 |
| P_T_ < 0.05 | -0.00015 | 0.00375 | 0.96785 |
| P_T_ < 0.1 | -0.00040 | 0.00394 | 0.91894 |
| P_T_ < 0.2 | 0.00020 | 0.00410 | 0.96059 |
| P_T_ < 0.3 | 0.00220 | 0.00419 | 0.59922 |
| P_T_ < 0.4 | 0.00202 | 0.00425 | 0.63497 |
| P_T_ < 0.5 | 0.00168 | 0.00428 | 0.69426 |
| P_T_ < 1 | 0.00174 | 0.00431 | 0.68666 |
| **Outcome: Subjective Well-Being Factor Score** | | | |
| **PRS Threshold** | **Beta** | **SE** | **P-val** |
| P_T_ < 5e.8 | -0.00032 | 0.00292 | 0.91281 |
| P_T_ < 1e.5 | 0.00042 | 0.00296 | 0.88633 |
| P_T_ < 0.001 | 0.00194 | 0.00311 | 0.53193 |
| P_T_ < 0.01 | 0.00044 | 0.00338 | 0.89700 |
| P_T_ < 0.05 | 0.00058 | 0.00375 | 0.87687 |
| P_T_ < 0.1 | 0.00068 | 0.00394 | 0.86314 |
| P_T_ < 0.2 | 0.00214 | 0.00410 | 0.60229 |
| P_T_ < 0.3 | 0.00277 | 0.00419 | 0.50892 |
| P_T_ < 0.4 | 0.00317 | 0.00425 | 0.45543 |
| P_T_ < 0.5 | 0.00294 | 0.00428 | 0.49238 |
| P_T_ < 1 | 0.00282 | 0.00430 | 0.51268 |
| **Outcome: Internalising Factor Score** | | | |
| **PRS Threshold** | **Beta** | **SE** | **P-val** |
| P_T_ < 5e.8 | 0.00055 | 0.00292 | 0.85201 |
| P_T_ < 1e.5 | 0.00074 | 0.00297 | 0.80204 |
| P_T_ < 0.001 | 0.00098 | 0.00311 | 0.75198 |
| P_T_ < 0.01 | -0.00133 | 0.00338 | 0.69422 |
| P_T_ < 0.05 | -0.00025 | 0.00375 | 0.94776 |
| P_T_ < 0.1 | -0.00069 | 0.00394 | 0.86041 |
| P_T_ < 0.2 | -0.00018 | 0.00410 | 0.96486 |
| P_T_ < 0.3 | 0.00164 | 0.00419 | 0.69550 |
| P_T_ < 0.4 | 0.00168 | 0.00425 | 0.69270 |
| P_T_ < 0.5 | 0.00140 | 0.00428 | 0.74348 |
| P_T_ < 1 | 0.00120 | 0.00430 | 0.78068 |

**Table S8.** Major depression polygenic risk score association test in case-only subsample

| **Outcome: Anxiety Factor Score** | | | |
| --- | --- | --- | --- |
| **PRS Threshold** | **Beta** | **SE** | **P-val** |
| P_T_ < 5e.8 | 0.01637 | 0.00673 | 0.01504 |
| P_T_ < 1e.5 | 0.00643 | 0.00672 | 0.33885 |
| P_T_ < 0.001 | 0.02142 | 0.00671 | 0.00141 |
| P_T_ < 0.01 | 0.04638 | 0.00670 | 4.56E-12 |
| P_T_ < 0.05 | 0.03824 | 0.00675 | 1.49E-08 |
| P_T_ < 0.1 | 0.04101 | 0.00678 | 1.44E-09 |
| P_T_ < 0.2 | 0.04375 | 0.00680 | 1.24E-10 |
| P_T_ < 0.3 | 0.04195 | 0.00681 | 7.54E-10 |
| P_T_ < 0.4 | 0.04213 | 0.00680 | 6.07E-10 |
| P_T_ < 0.5 | 0.04168 | 0.00680 | 8.74E-10 |
| P_T_ < 1 | 0.04062 | 0.00680 | 2.30E-09 |
| **Outcome: Mood Factor Score** | | | |
| **PRS Threshold** | **Beta** | **SE** | **P-val** |
| P_T_ < 5e.8 | 0.01119 | 0.00659 | 0.08968 |
| P_T_ < 1e.5 | 0.00950 | 0.00658 | 0.14846 |
| P_T_ < 0.001 | 0.02242 | 0.00657 | 0.00064 |
| P_T_ < 0.01 | 0.04576 | 0.00656 | 3.09E-12 |
| P_T_ < 0.05 | 0.04677 | 0.00661 | 1.48E-12 |
| P_T_ < 0.1 | 0.05057 | 0.00663 | 2.47E-14 |
| P_T_ < 0.2 | 0.05464 | 0.00665 | 2.21E-16 |
| P_T_ < 0.3 | 0.05419 | 0.00667 | 4.55E-16 |
| P_T_ < 0.4 | 0.05375 | 0.00666 | 7.19E-16 |
| P_T_ < 0.5 | 0.05307 | 0.00665 | 1.51E-15 |
| P_T_ < 1 | 0.05366 | 0.00665 | 7.40E-16 |
| **Outcome: Neurovegetative Factor Score** | | | |
| **PRS Threshold** | **Beta** | **SE** | **P-val** |
| P_T_ < 5e.8 | 0.01182 | 0.00657 | 0.07191 |
| P_T_ < 1e.5 | 0.00462 | 0.00655 | 0.48098 |
| P_T_ < 0.001 | 0.02223 | 0.00654 | 0.00068 |
| P_T_ < 0.01 | 0.04317 | 0.00653 | 3.96E-11 |
| P_T_ < 0.05 | 0.04745 | 0.00658 | 5.65E-13 |
| P_T_ < 0.1 | 0.05215 | 0.00660 | 2.92E-15 |
| P_T_ < 0.2 | 0.05625 | 0.00662 | 2.10E-17 |
| P_T_ < 0.3 | 0.05526 | 0.00664 | 8.95E-17 |
| P_T_ < 0.4 | 0.05477 | 0.00663 | 1.53E-16 |
| P_T_ < 0.5 | 0.05400 | 0.00662 | 3.65E-16 |
| P_T_ < 1 | 0.05412 | 0.00662 | 3.15E-16 |

| **Outcome: Psychomotor Cognitive Factor Score** | | | |
| --- | --- | --- | --- |
| **PRS Threshold** | **Beta** | **SE** | **P-val** |
| P_T_ < 5e.8 | 0.01312 | 0.00667 | 0.04920 |
| P_T_ < 1e.5 | 0.00800 | 0.00665 | 0.22908 |
| P_T_ < 0.001 | 0.02551 | 0.00665 | 0.00012 |
| P_T_ < 0.01 | 0.04817 | 0.00664 | 3.99E-13 |
| P_T_ < 0.05 | 0.04909 | 0.00668 | 2.11E-13 |
| P_T_ < 0.1 | 0.05265 | 0.00671 | 4.32E-15 |
| P_T_ < 0.2 | 0.05528 | 0.00673 | 2.19E-16 |
| P_T_ < 0.3 | 0.05402 | 0.00675 | 1.21E-15 |
| P_T_ < 0.4 | 0.05333 | 0.00674 | 2.53E-15 |
| P_T_ < 0.5 | 0.05261 | 0.00673 | 5.48E-15 |
| P_T_ < 1 | 0.05248 | 0.00673 | 6.42E-15 |
| **Outcome: Subjective Well-Being Factor Score** | | | |
| **PRS Threshold** | **Beta** | **SE** | **P-val** |
| P_T_ < 5e.8 | 0.00658 | 0.00647 | 0.30934 |
| P_T_ < 1e.5 | 0.00933 | 0.00645 | 0.14833 |
| P_T_ < 0.001 | 0.02038 | 0.00645 | 0.00157 |
| P_T_ < 0.01 | 0.03985 | 0.00644 | 6.07E-10 |
| P_T_ < 0.05 | 0.03973 | 0.00648 | 9.04E-10 |
| P_T_ < 0.1 | 0.04141 | 0.00651 | 2.00E-10 |
| P_T_ < 0.2 | 0.04695 | 0.00653 | 6.52E-13 |
| P_T_ < 0.3 | 0.04757 | 0.00654 | 3.73E-13 |
| P_T_ < 0.4 | 0.04641 | 0.00654 | 1.26E-12 |
| P_T_ < 0.5 | 0.04604 | 0.00653 | 1.78E-12 |
| P_T_ < 1 | 0.04650 | 0.00653 | 1.06E-12 |
| **Outcome: Internalising Factor Score** | | | |
| **PRS Threshold** | **Beta** | **SE** | **P-val** |
| P_T_ < 5e.8 | 0.01215 | 0.00658 | 0.06508 |
| P_T_ < 1e.5 | 0.00853 | 0.00657 | 0.19394 |
| P_T_ < 0.001 | 0.02350 | 0.00656 | 0.00034 |
| P_T_ < 0.01 | 0.04704 | 0.00655 | 7.07E-13 |
| P_T_ < 0.05 | 0.04771 | 0.00660 | 4.90E-13 |
| P_T_ < 0.1 | 0.05150 | 0.00662 | 7.59E-15 |
| P_T_ < 0.2 | 0.05546 | 0.00664 | 7.18E-17 |
| P_T_ < 0.3 | 0.05472 | 0.00666 | 2.15E-16 |
| P_T_ < 0.4 | 0.05421 | 0.00665 | 3.70E-16 |
| P_T_ < 0.5 | 0.05353 | 0.00664 | 7.87E-16 |
| P_T_ < 1 | 0.05379 | 0.00664 | 5.76E-16 |

**Table S9.** Major depression polygenic risk score association test in control-only subsample

| **Outcome: Anxiety Factor Score** | | | |
| --- | --- | --- | --- |
| **PRS Threshold** | **Beta** | **SE** | **P-val** |
| P_T_ < 5e.8 | 0.00164 | 0.00299 | 0.58212 |
| P_T_ < 1e.5 | 0.00947 | 0.00298 | 0.00151 |
| P_T_ < 0.001 | 0.01173 | 0.00299 | 8.97E-05 |
| P_T_ < 0.01 | 0.01838 | 0.00301 | 9.93E-10 |
| P_T_ < 0.05 | 0.01869 | 0.00301 | 5.61E-10 |
| P_T_ < 0.1 | 0.02019 | 0.00303 | 2.64E-11 |
| P_T_ < 0.2 | 0.02143 | 0.00304 | 1.69E-12 |
| P_T_ < 0.3 | 0.02232 | 0.00303 | 1.89E-13 |
| P_T_ < 0.4 | 0.02185 | 0.00303 | 6.16E-13 |
| P_T_ < 0.5 | 0.02183 | 0.00303 | 6.07E-13 |
| P_T_ < 1 | 0.02205 | 0.00303 | 3.63E-13 |
| **Outcome: Mood Factor Score** | | | |
| **PRS Threshold** | **Beta** | **SE** | **P-val** |
| P_T_ < 5e.8 | -0.00008 | 0.00283 | 0.97727 |
| P_T_ < 1e.5 | 0.00644 | 0.00283 | 0.02290 |
| P_T_ < 0.001 | 0.01165 | 0.00284 | 4.04E-05 |
| P_T_ < 0.01 | 0.01966 | 0.00285 | 5.37E-12 |
| P_T_ < 0.05 | 0.02097 | 0.00286 | 2.14E-13 |
| P_T_ < 0.1 | 0.02234 | 0.00287 | 7.04E-15 |
| P_T_ < 0.2 | 0.02299 | 0.00288 | 1.34E-15 |
| P_T_ < 0.3 | 0.02367 | 0.00288 | 1.86E-16 |
| P_T_ < 0.4 | 0.02322 | 0.00288 | 6.97E-16 |
| P_T_ < 0.5 | 0.02347 | 0.00287 | 3.21E-16 |
| P_T_ < 1 | 0.02380 | 0.00287 | 1.25E-16 |
| **Outcome: Neurovegetative Factor Score** | | | |
| **PRS Threshold** | **Beta** | **SE** | **P-val** |
| P_T_ < 5e.8 | -0.00050 | 0.00285 | 0.86149 |
| P_T_ < 1e.5 | 0.00496 | 0.00285 | 0.08121 |
| P_T_ < 0.001 | 0.01076 | 0.00286 | 0.00016 |
| P_T_ < 0.01 | 0.02046 | 0.00287 | 9.94E-13 |
| P_T_ < 0.05 | 0.02215 | 0.00287 | 1.31E-14 |
| P_T_ < 0.1 | 0.02356 | 0.00289 | 3.46E-16 |
| P_T_ < 0.2 | 0.02427 | 0.00289 | 5.20E-17 |
| P_T_ < 0.3 | 0.02469 | 0.00289 | 1.43E-17 |
| P_T_ < 0.4 | 0.02434 | 0.00289 | 4.17E-17 |
| P_T_ < 0.5 | 0.02462 | 0.00289 | 1.74E-17 |
| P_T_ < 1 | 0.02514 | 0.00289 | 3.61E-18 |

| **Outcome: Psychomotor Cognitive Factor Score** | | | |
| --- | --- | --- | --- |
| **PRS Threshold** | **Beta** | **SE** | **P-val** |
| P_T_ < 5e.8 | -0.00017 | 0.00280 | 0.95120 |
| P_T_ < 1e.5 | 0.00598 | 0.00280 | 0.03255 |
| P_T_ < 0.001 | 0.01071 | 0.00280 | 0.00013 |
| P_T_ < 0.01 | 0.01917 | 0.00282 | 1.03E-11 |
| P_T_ < 0.05 | 0.01975 | 0.00282 | 2.67E-12 |
| P_T_ < 0.1 | 0.02114 | 0.00284 | 9.09E-14 |
| P_T_ < 0.2 | 0.02199 | 0.00284 | 1.05E-14 |
| P_T_ < 0.3 | 0.02273 | 0.00284 | 1.28E-15 |
| P_T_ < 0.4 | 0.02253 | 0.00284 | 2.28E-15 |
| P_T_ < 0.5 | 0.02277 | 0.00284 | 1.11E-15 |
| P_T_ < 1 | 0.02328 | 0.00284 | 2.56E-16 |
| **Outcome: Subjective Well-Being Factor Score** | | | |
| **PRS Threshold** | **Beta** | **SE** | **P-val** |
| P_T_ < 5e.8 | -0.00481 | 0.00330 | 0.14584 |
| P_T_ < 1e.5 | 0.00632 | 0.00330 | 0.05560 |
| P_T_ < 0.001 | 0.01228 | 0.00331 | 0.00021 |
| P_T_ < 0.01 | 0.01979 | 0.00333 | 2.77E-09 |
| P_T_ < 0.05 | 0.02070 | 0.00333 | 5.47E-10 |
| P_T_ < 0.1 | 0.02247 | 0.00335 | 2.00E-11 |
| P_T_ < 0.2 | 0.02466 | 0.00336 | 2.11E-13 |
| P_T_ < 0.3 | 0.02517 | 0.00336 | 6.60E-14 |
| P_T_ < 0.4 | 0.02482 | 0.00336 | 1.46E-13 |
| P_T_ < 0.5 | 0.02528 | 0.00336 | 4.99E-14 |
| P_T_ < 1 | 0.02569 | 0.00336 | 1.98E-14 |
| **Outcome: Internalising Factor Score** | | | |
| **PRS Threshold** | **Beta** | **SE** | **P-val** |
| P_T_ < 5e.8 | -0.00035 | 0.00284 | 0.90086 |
| P_T_ < 1e.5 | 0.00661 | 0.00284 | 0.01999 |
| P_T_ < 0.001 | 0.01181 | 0.00285 | 3.39E-05 |
| P_T_ < 0.01 | 0.02029 | 0.00286 | 1.33E-12 |
| P_T_ < 0.05 | 0.02146 | 0.00287 | 7.18E-14 |
| P_T_ < 0.1 | 0.02294 | 0.00288 | 1.72E-15 |
| P_T_ < 0.2 | 0.02382 | 0.00289 | 1.65E-16 |
| P_T_ < 0.3 | 0.02450 | 0.00289 | 2.16E-17 |
| P_T_ < 0.4 | 0.02409 | 0.00289 | 7.25E-17 |
| P_T_ < 0.5 | 0.02435 | 0.00288 | 3.25E-17 |
| P_T_ < 1 | 0.02475 | 0.00289 | 9.94E-18 |

**Table S10.** Major depression polygenic risk score association test in unknown-only subsample

| **Outcome: Anxiety Factor Score** | | | |
| --- | --- | --- | --- |
| **PRS Threshold** | **Beta** | **SE** | **P-val** |
| P_T_ < 5e.8 | 0.00847 | 0.00572 | 0.13889 |
| P_T_ < 1e.5 | 0.00665 | 0.00577 | 0.24925 |
| P_T_ < 0.001 | 0.00986 | 0.00576 | 0.08662 |
| P_T_ < 0.01 | 0.01679 | 0.00576 | 0.00357 |
| P_T_ < 0.05 | 0.02510 | 0.00576 | 1.34E-05 |
| P_T_ < 0.1 | 0.02581 | 0.00579 | 8.19E-06 |
| P_T_ < 0.2 | 0.02427 | 0.00581 | 3.00E-05 |
| P_T_ < 0.3 | 0.02404 | 0.00582 | 3.58E-05 |
| P_T_ < 0.4 | 0.02102 | 0.00583 | 0.00031 |
| P_T_ < 0.5 | 0.02106 | 0.00583 | 0.00030 |
| P_T_ < 1 | 0.02180 | 0.00583 | 0.00019 |
| **Outcome: Mood Factor Score** | | | |
| **PRS Threshold** | **Beta** | **SE** | **P-val** |
| P_T_ < 5e.8 | 0.01384 | 0.00572 | 0.01555 |
| P_T_ < 1e.5 | 0.00846 | 0.00577 | 0.14259 |
| P_T_ < 0.001 | 0.01340 | 0.00575 | 0.01985 |
| P_T_ < 0.01 | 0.01859 | 0.00576 | 0.00124 |
| P_T_ < 0.05 | 0.02601 | 0.00576 | 6.35E-06 |
| P_T_ < 0.1 | 0.02555 | 0.00578 | 9.97E-06 |
| P_T_ < 0.2 | 0.02689 | 0.00581 | 3.71E-06 |
| P_T_ < 0.3 | 0.02653 | 0.00581 | 5.04E-06 |
| P_T_ < 0.4 | 0.02369 | 0.00582 | 4.76E-05 |
| P_T_ < 0.5 | 0.02467 | 0.00583 | 2.30E-05 |
| P_T_ < 1 | 0.02511 | 0.00583 | 1.67E-05 |
| **Outcome: Neurovegetative Factor Score** | | | |
| **PRS Threshold** | **Beta** | **SE** | **P-val** |
| P_T_ < 5e.8 | 0.01478 | 0.00567 | 0.00909 |
| P_T_ < 1e.5 | 0.00406 | 0.00572 | 0.47725 |
| P_T_ < 0.001 | 0.00881 | 0.00570 | 0.12245 |
| P_T_ < 0.01 | 0.01806 | 0.00571 | 0.00155 |
| P_T_ < 0.05 | 0.02479 | 0.00571 | 1.41E-05 |
| P_T_ < 0.1 | 0.02508 | 0.00573 | 1.22E-05 |
| P_T_ < 0.2 | 0.02396 | 0.00576 | 3.19E-05 |
| P_T_ < 0.3 | 0.02399 | 0.00576 | 3.14E-05 |
| P_T_ < 0.4 | 0.02171 | 0.00577 | 0.00017 |
| P_T_ < 0.5 | 0.02296 | 0.00577 | 7.03E-05 |
| P_T_ < 1 | 0.02318 | 0.00578 | 6.07E-05 |

| **Outcome: Psychomotor Cognitive Factor Score** | | | |
| --- | --- | --- | --- |
| **PRS Threshold** | **Beta** | **SE** | **P-val** |
| P_T_ < 5e.8 | 0.01547 | 0.00572 | 0.00684 |
| P_T_ < 1e.5 | 0.01071 | 0.00577 | 0.06345 |
| P_T_ < 0.001 | 0.01613 | 0.00575 | 0.00504 |
| P_T_ < 0.01 | 0.02047 | 0.00576 | 0.00038 |
| P_T_ < 0.05 | 0.02897 | 0.00576 | 4.96E-07 |
| P_T_ < 0.1 | 0.02843 | 0.00578 | 8.86E-07 |
| P_T_ < 0.2 | 0.02813 | 0.00581 | 1.29E-06 |
| P_T_ < 0.3 | 0.02865 | 0.00581 | 8.33E-07 |
| P_T_ < 0.4 | 0.02630 | 0.00582 | 6.35E-06 |
| P_T_ < 0.5 | 0.02703 | 0.00583 | 3.50E-06 |
| P_T_ < 1 | 0.02740 | 0.00583 | 2.62E-06 |
| **Outcome: Subjective Well-Being Factor Score** | | | |
| **PRS Threshold** | **Beta** | **SE** | **P-val** |
| P_T_ < 5e.8 | 0.01322 | 0.00564 | 0.01903 |
| P_T_ < 1e.5 | 0.01315 | 0.00569 | 0.02078 |
| P_T_ < 0.001 | 0.01429 | 0.00567 | 0.01172 |
| P_T_ < 0.01 | 0.01963 | 0.00568 | 0.00054 |
| P_T_ < 0.05 | 0.02911 | 0.00568 | 2.97E-07 |
| P_T_ < 0.1 | 0.02894 | 0.00570 | 3.89E-07 |
| P_T_ < 0.2 | 0.02958 | 0.00573 | 2.42E-07 |
| P_T_ < 0.3 | 0.02914 | 0.00573 | 3.72E-07 |
| P_T_ < 0.4 | 0.02764 | 0.00574 | 1.49E-06 |
| P_T_ < 0.5 | 0.02797 | 0.00574 | 1.13E-06 |
| P_T_ < 1 | 0.02844 | 0.00575 | 7.56E-07 |
| **Outcome: Internalising Factor Score** | | | |
| **PRS Threshold** | **Beta** | **SE** | **P-val** |
| P_T_ < 5e.8 | 0.01426 | 0.00564 | 0.01149 |
| P_T_ < 1e.5 | 0.00865 | 0.00569 | 0.12840 |
| P_T_ < 0.001 | 0.01340 | 0.00567 | 0.01818 |
| P_T_ < 0.01 | 0.01942 | 0.00568 | 0.00063 |
| P_T_ < 0.05 | 0.02741 | 0.00568 | 1.41E-06 |
| P_T_ < 0.1 | 0.02716 | 0.00570 | 1.93E-06 |
| P_T_ < 0.2 | 0.02757 | 0.00573 | 1.51E-06 |
| P_T_ < 0.3 | 0.02743 | 0.00573 | 1.74E-06 |
| P_T_ < 0.4 | 0.02474 | 0.00574 | 1.67E-05 |
| P_T_ < 0.5 | 0.02562 | 0.00575 | 8.27E-06 |
| P_T_ < 1 | 0.02606 | 0.00575 | 5.89E-06 |

**Supplementary References**

Bycroft, C., Freeman, C., Petkova, D., Band, G., Elliott, L.T., Sharp, K., … Marchini, J. (2018). The UK Biobank resource with deep phenotyping and genomic data. *Nature, 562*(7726), 203. doi: <10.1038/s41586-018-0579-z>

Manichaikul, A., Mychaleckyj, J.C., Rich, S.S., Daly, K., Sale, M., Chen, W.M. (2010) Robust relationship inference in genome-wide association studies. *Bioinformatics, 26*(22), 2867-73. doi: [10.1093/bioinformatics/btq559](https://doi.org/10.1093/bioinformatics/btq559)

Warren, H.R., Evangelou, E., Cabrera, C.P., Gao, H., Ren, M., Mifsud, B. ... UK Biobank CardioMetabolic Consortium BP working group. (2017). Genome-wide association analysis identifies novel blood pressure loci and offers biological insights into cardiovascular risk. *Nature genetics, 49*(3), 403– 415. doi: [10.1038/ng.3768](https://doi.org/10.1038/ng.3768)

Abraham, G., Qiu, Y., Inouye, M. (2017). FlashPCA2: principal component analysis of Biobank-scale genotype datasets. *Bioinformatics, 33*, 2776–2778. doi: [10.1093/bioinformatics/btx299](https://doi.org/10.1093/bioinformatics/btx299)

Davis, K. A., Coleman, J. R., Adams, M., Allen, N., Breen, G., Cullen, B., ... Hotopf, M. (2020). Mental health in UK Biobank–development, implementation and results from an online questionnaire completed by 157 366 participants: a reanalysis. *BJPsych open*, *6*(2). doi: [10.1192/bjo.2019.100](https://doi.org/10.1192/bjo.2019.100)

Coleman, J. R., Peyrot, W. J., Purves, K. L., Davis, K. A., Rayner, C., Choi, S. W., ... Breen, G. (2020). Genome-wide gene-environment analyses of major depressive disorder and reported lifetime traumatic experiences in UK Biobank. *Molecular psychiatry*, 1-17. doi: <10.1038/s41380-019-0546-6>
